# Enhancer-priming in ageing human bone marrow mesenchymal stromal cells contributes to immune traits

**DOI:** 10.1101/2021.09.03.458728

**Authors:** Mang Ching Lai, Mariana Ruiz-Velasco, Christian Arnold, Olga Sigalova, Daria Bunina, Ivan Berest, Ximing Ding, Marco L. Hennrich, Laura Poisa-Beiro, Annique Claringbould, Anna Mathioudaki, Caroline Pabst, Anthony D. Ho, Anne-Claude Gavin, Judith B. Zaugg

## Abstract

Bone marrow mesenchymal stromal cells (BMSCs) can differentiate into adipocytes and osteoblasts, and are important regulators of the haematopoietic system. Ageing associates with an increased ratio of bone marrow adipocytes to osteoblasts and immune dysregulation. Here, we carried out an integrative multiomics analysis of ATAC-Seq, RNA-Seq and proteomics data from primary human BMSCs in a healthy cohort age between 20 - 60. We identified age-sensitive elements uniquely affecting each molecular level where transcription is mostly spared, and characterised the underlying biological pathways, revealing the interplay of age-related gene expression mechanism changes spanning multiple gene regulatory layers. Through data integration with enhancer-mediated gene regulatory network analysis, we discovered that enhancers and transcription factors influence cell differentiation potential in the ageing BMSCs. By combining our results with genome-wide association study data, we found that age-specific changes could contribute to common traits related to BMSC-derived tissues such as bone and adipose tissue, and to immune-related traits on a systemic level such as asthma. We demonstrate here that a multiomics approach is crucial for unravelling complex information, providing new insights on how ageing contributes to bone marrow- and immune-related disorders.

## Introduction

Ageing in humans is characterised by molecular alterations that together affect cellular functions, ultimately contributing to functional decline at the organism level (López-Otín et al., 2013). The ageing of stem cells in particular is a fundamental mechanism in causing undesirable age-related changes in tissues and organs since stem cells could accumulate a lifetime of environmental, genetic and epigenetic alterations, and the effects amplified at the organism level through clonal expansion (Ermolaeva et al., 2018). The age-related exhaustion of stem cells, where stem cells lose their differentiation and self-renewal potential, further exacerbates the ageing phenotype of an organism (López-Otín et al., 2013). However, the molecular mechanisms remain largely elusive.

The bone marrow (BM) harbours two important stem and progenitor cell types, haematopoietic stem and progenitor cells (HSPCs) and mesenchymal stem cells (MSCs). MSCs represent a minor cell population, an estimated <0.1% of mononuclear cells (Nombela-Arrieta and Manz, 2017) in the bone marrow, and are the precursors of osteoblasts, chondrocytes and adipocytes (Friedman et al., 2006; Short et al., 2003). MSCs are described as critical regulators of hematopoietic cells and crucial for HSPC function (Morrison and Scadden, 2014; Yu and Scadden, 2016). For example, they keep HSPCs in a quiescent stem-cell state *in vivo* (Gottschling et al., 2007) and *in vitro* (Jing et al., 2010). MSCs have also demonstrated immunomodulatory roles, potentially regulating immune cell functions via both cell-cell interaction and secreted cytokines (Gao et al., 2016). Due to their low abundance, human MSCs are mostly studied in *in vitro* models where primary BM-derived mesenchymal stem/stromal cells (BMSCs) can be expanded in culture. Cultured BMSCs keep their multipotent differentiation potential, and are thus a good model system to investigate MSC properties.

During ageing, human and mouse BMSCs have been described to lose both their general function and differentiation potential (Kretlow et al., 2008; Mueller and Glowacki, 2001; Yin et al., 2017), with the ratio of adipocytes to osteoblasts in the bone marrow increasing with age (Meunier et al., 1971; Moerman et al., 2004; Short et al., 2003). The age-related adipogenic shift has recently been further demonstrated at single cell resolution in mMSCs (Dolgalev and Tikhonova, 2021; Woods and Guezguez, 2021; Zhong et al., 2020). The differentiation of MSC/BMSCs into mature cell types is mostly driven by a specific set of transcription factors (TFs) including CEBPs, RUNX2, FOXO and PARP (Chen et al., 2016). TFs driving BMSC differentiation into osteoblasts and adipocytes *in vitro* have been shown to regulate a highly interconnected network of enhancers to drive BMSCs differentiation into osteoblasts and adipocytes *in vitro (Chen et al., 2016; Rauch et al., 2019)*. TFs-enhancer networks therefore represent a complex layer of information, inaccessible by individual genomic profiling methods, on the potential of the cells to respond to cues during differentiation or perturbation, which only becomes visible when integrating complementary omics approaches. While our previous study uncovered the age-dependent synergistic misregulation of BMSC function in the central carbon and glycolytic pathways in the cell proteome (Hennrich et al., 2018), the mechanisms of how ageing affects the intricate TF-enhancer network and therefore cell potential is not understood.

The goal of this study is to jointly interrogate cellular function and differentiation potential in human BMSCs during ageing to understand the molecular and cellular mechanisms that may drive ageing-related changes in the bone marrow niche, and to explore how these changes may contribute to age-dependent common traits and diseases. We performed an integrative multiomics analysis in BMSCs in a cohort of healthy individuals between 20 and 60 years old, combining chromatin accessibility profiling (ATAC-seq) with RNA expression (Pellagatti et al., 2018) and protein abundance measured by mass spectrometry (Hennrich et al., 2018). Our results suggest that ageing simultaneously impairs cellular function via post-transcriptional gene expression mechanisms, and cell differentiation potential via chromatin priming. Using enhancer-mediated gene regulatory networks, we discovered that age-dependent priming of BMSC enhancers could lead to adipogenic bias via synergistic TF activity changes acting on their target gene. By further combining the BMSC adipogenic network with genome-wide association study (GWAS) data, we identified age-sensitive TFs that could contribute to immune-related GWAS traits on a systemic level. Here we showcase an integrative strategy, by combining BMSC datasets together with publicly available databases, that is necessary to uncover the complex layers of age-related effects on cell potential and function.

## Results

### Multi-omics data from ageing bone marrow-derived mesenchymal stromal cells

To study the molecular mechanisms of age-specific changes in the human bone marrow-derived mesenchymal stromal cells (BMSCs), we assembled a dataset comprising of ATAC-Seq, RNA-Seq and proteomics data, generated from BMSCs obtained from a cohort of healthy donors between 20 to 60 years old, representing snapshots of the “ageing” process (**Figure 1A**). Briefly, BMSCs were obtained by culturing mononuclear cells obtained from bone marrow aspirates and harvested at passage four for the different molecular assays (see Methods). The mass spectrometry and the RNA-Seq data were previously generated (Hennrich et al., 2018; Pellagatti et al., 2018), and were reanalysed in this study (supplementary material, **Figure S1D-E**) while chromatin accessibility data from the same cohort of donors was generated in this study (see Methods).

**Figure 1.**
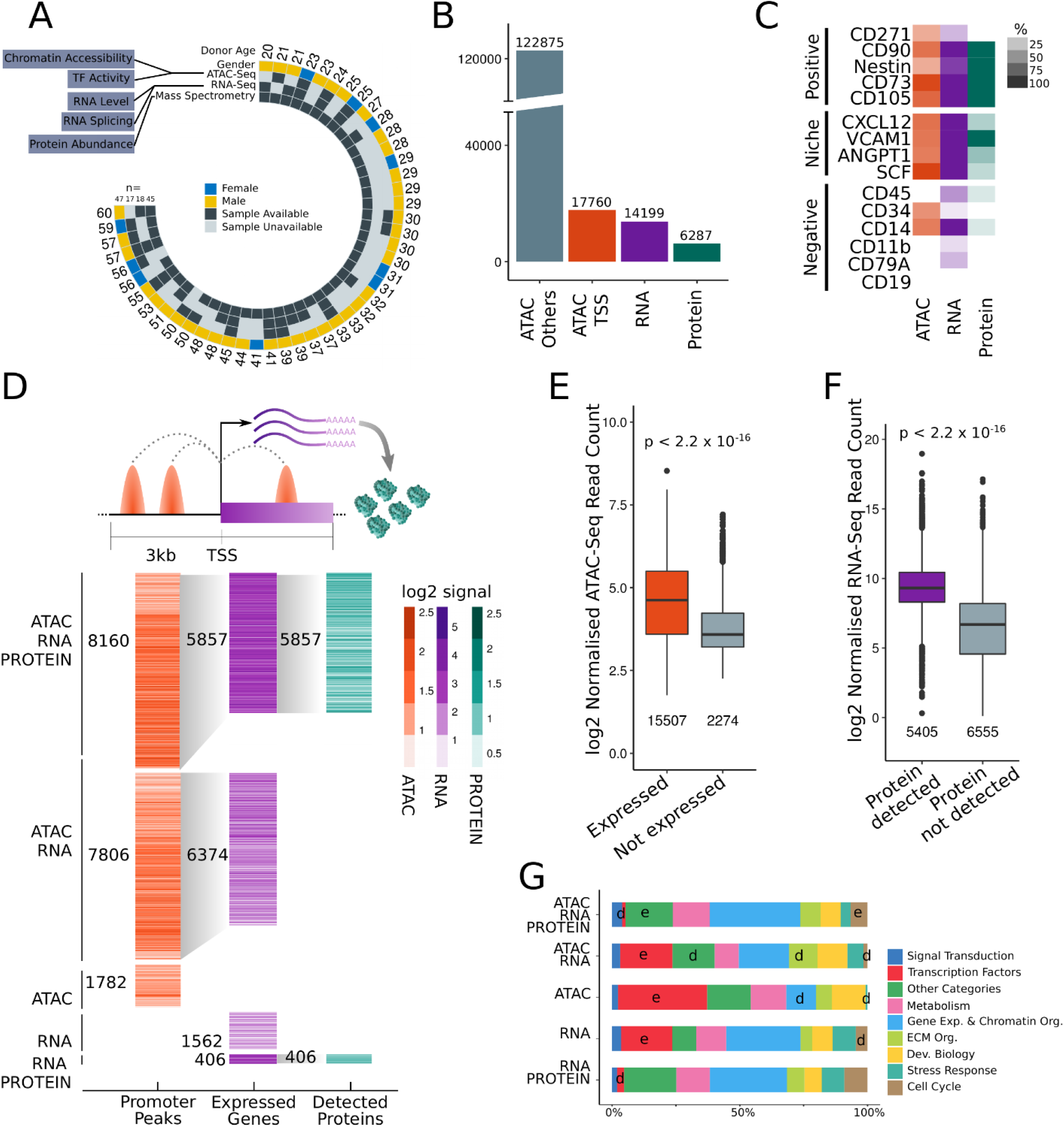
Multiomics profiling in bone marrow-derived mesenchymal stromal cells (BMSCs) across an ageing cohort of healthy individuals. **A)** Overview of the samples used in this study for ATAC-Seq, RNA-Seq and mass spectrometry analysis. **B)** Number of peaks, genes and proteins measured by ATAC-Seq, RNA-Seq and mass spectrometry. ATAC-Seq peaks are further classified into promoter peaks near transcription start (+/-3kb) and other peaks. **C)** Positive and negative BMSC marker genes and niche factor genes measured in BMSCs by different assays are shown as the percentage of individuals in which the marker was detected. The log2-transformed signal of the normalised read counts (RNA-seq ATAC-seq) or protein abundances are shown for each gene captured by ATAC-promoter peaks, RNA-Seq or mass spectrometry in BMSCs. Numbers indicate the unique features in each category. Note that a gene can have multiple promoter ATAC-seq peaks. **E)** Distributions of log2-transformed normalised read count of ATAC-Seq promoter peaks are shown for all genes stratified by whether or not they are detected in RNA-Seq data (p-value from Wilcox’s test). **F)** Distributions of log2-transformed normalised read count of RNA-Seq transcripts are shown for genes stratified by whether or not the gene was detected in proteomics data (p-value from Wilcox’s test). **G)** Percentage of genes in different reactome pathway categories, captured by ATAC-Seq, RNA-Seq and proteomics data. Enrichment e, depletion d (Fisher’s exact test, p-adjust < 0.1).

In total, we profiled BMSCs using ATAC-Seq in 17 individuals with two replicated cell cultures, and obtained RNA-Seq and proteomics data from 18 and 45 donors, respectively. The ATAC-seq data was processed using our in-house pipeline, including mapping, peak-calling and quality controls (see Methods). Only samples that passed all standard quality controls were used for further analysis. We defined a consensus peakset of 140,635 peaks that are present in at least 4 of the 34 samples, equivalent to at least 2 individuals (**Figure S1A**). We defined promoter peaks as ATAC-Seq peaks that are within 3 kb of the transcription start site (TSS) of a protein coding gene. Promoter peaks account for about 14.5 % of all ATAC-Seq peaks. This resulted in four types of quantified features: genome-wide ATAC-seq peaks, promoter ATAC-seq peaks, RNA and protein abundances (**Figure 1B**).

We first examined the presence of BMSC positive (CD271, CD90, Nestin, CD73, CD105), and negative markers (CD45, CD34, CD14, CD11b, and CD79a), as well as secreted niche factors (CXCL12, VCAM1, ANGPT1, SCF) (Calvi and Link, 2015; Dominici et al., 2006; Nakahara et al., 2019) in the promoter ATAC-Seq, RNA-Seq and proteomics data. The presence of all the positive markers and niche factors, and the absence of the majority of the negative markers, confirms the stromal identity of our cells across the three assays (**Figure 1C**). We further compared the BMSC ATAC-Seq profiles with previously published BMSC DNase-Seq profiles (Roadmap Epigenomics Consortium et al., 2015) to ensure that the accessible regions share a similar global genomic distribution (**Figure S1B-C**). The BMSC-specific expression and accessibility profiles indicate that the cell population we obtained after four passages of culturing *in vitro* retains *bona fide* mesenchymal multipotent stromal cell characteristics.

To obtain a gene-centric global overview across the three omics assays, we assessed which genes are quantified by promoter ATAC-Seq peak accessibility, RNA level and protein abundance. Approximately 90% (15,966 peaks) of the genes with an accessible promoter peak are quantified by RNA-Seq and 33% (5,857 genes) by mass spectrometry. Of the 14,199 genes quantified by RNA-Seq, 44% (6287 genes) are captured by mass spectrometry. About 86% of the genes measured by RNA-Seq (12,231) and 93% of the quantified BMSC proteome (5,857) contained an ATAC-seq peak in their promoter region (**Figure 1D**).

We found that the promoter ATAC-Seq peaks are significantly more accessible when their associated gene was measured by RNA-seq (Wilcox test, p < 2.2 × 10^−16^) (**Figure 1E**). Similarly, the RNA-seq read counts showed a significantly higher level when their corresponding proteins are detected by mass spectrometry (Wilcox test, p < 2.2 × 10^−16^) (**Figure 1F**). The different levels of gene signals detected by RNA-seq vs mass spectrometry likely reflects the difference in sensitivity of the methods, and also post-transcriptional and translational regulation. The promoter regions where we detect an ATAC-Seq peak but no expression by RNA-seq, may reflect poised promoters whose accessibility precedes the transcription of the gene in a differentiation process as also observed on other stem cells (Bunina et al., 2020).

We found that genes captured by all three profiling methods, ATAC-Seq promoter peaks, RNA-Seq and mass spectrometry, are depleted for being sequence-specific TFs while being enriched in cell cycle, extracellular matrix organisation and other pathways such as transmembrane transport of small molecules (**Figure 1G**). On the other hand, genes captured by ATAC-Seq promoter peaks and/or RNA-Seq only, are significantly enriched for being TFs. The results show that sequence-specific TFs are systematically missed in mass spectrometry measurements relative to ATAC-Seq and RNA-Seq assays. This observation is in agreement with previous evidence that TFs are low abundant proteins that are often difficult to capture by proteomics approaches (Ding et al., 2013; Tacheny et al., 2012).

The different global profiling methodologies provided different coverage of the cellular components, suggesting that specific gene categories, such as TFs, are understudied when using proteomics data only. The results here showed the importance of capturing the epigenomic landscape using ATAC-Seq to complement RNA-Seq and proteomics methodologies.

### Ageing hotspots in BMSCs affect multiple mechanisms regulating gene expression

Using the ATAC-Seq, RNA-Seq and proteomics data, we quantified the impact of ageing on five different molecular levels: From ATAC-seq data, we obtained chromatin accessibility and TF activity; from RNA-seq data we obtained RNA expression and differentially used exon levels; from the proteomics data we obtained protein abundance (**Figure 1A**, see Methods). To quantify age-related changes in the BMSC chromatin accessibility, we performed differential peak analysis using DESeq2 (Love et al., 2014) (see Methods). We identified 3,852 differentially accessible peaks at 10% false discovery rate (FDR) from the consensus peakset of 140,635 peaks, of which 2,578 are ageing-down and 1,274 ageing-up (**Figure 2A-B**). Mapping these age-sensitive ATAC-seq peaks to gene promoters (+/3kb from the TSS) revealed 659 genes with an age-sensitive promoter.

**Figure 2.**
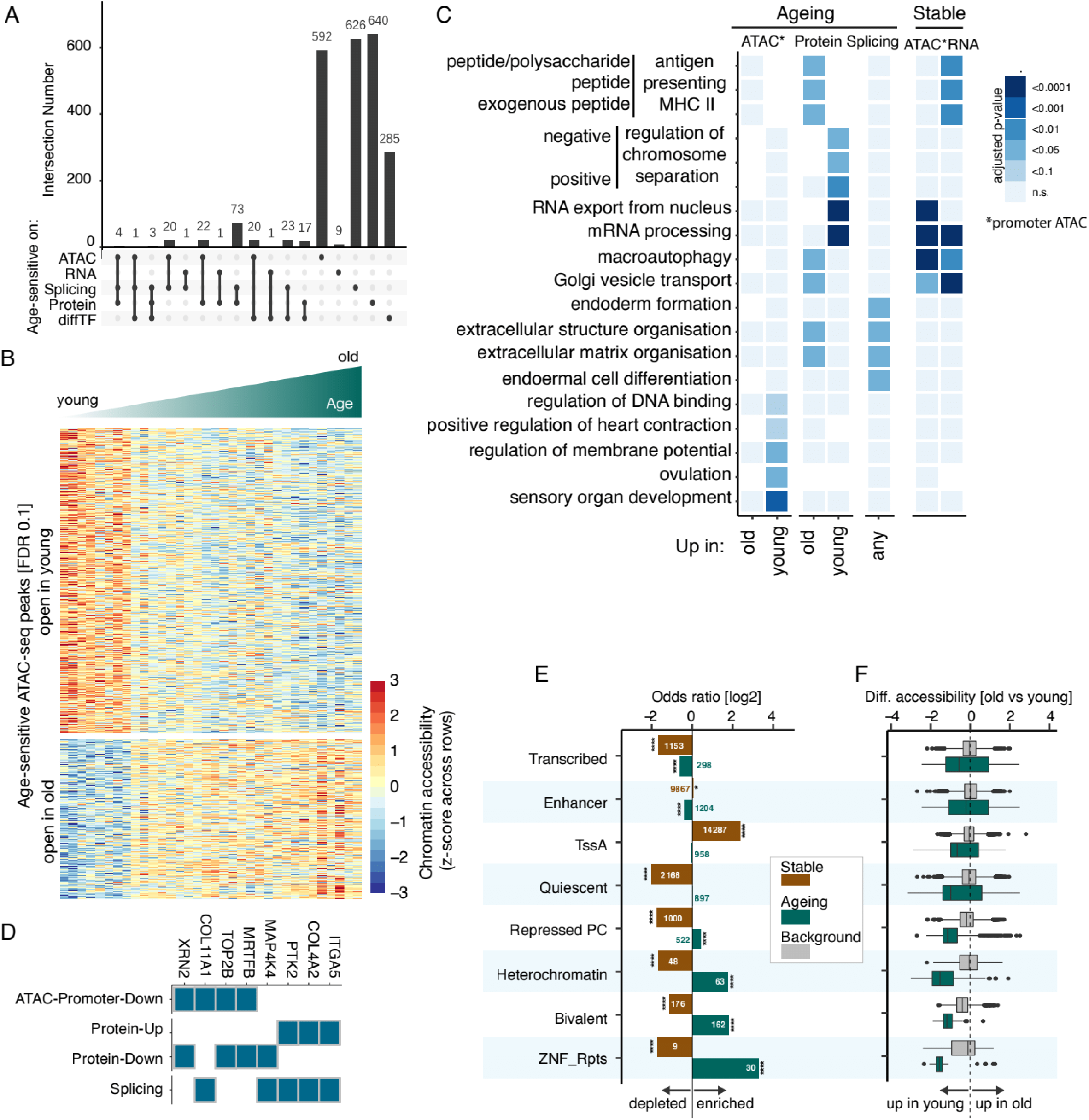
Age-hotspots exist on chromatin accessibility, splicing, protein expression and transcription factor activity. **A)** The intersect number of age-sensitive genes that are age-sensitive on the different molecular levels. **B)** Heatmap showing the row z-score of the normalised ATAC-Seq peak read counts for all age-sensitive peaks. Columns represent samples and are ordered from young to old, and rows represent ATAC-Seq peak stratified by whether they are open in young or old. **C)** Enrichment of GO terms of genes that are age-sensitive on the different molecular levels (ATAC: promoter accessibility, Protein: abundance, splicing: exon usage). **D)** Genes in the enriched GO terms that are age-sensitive on multiple layers are shown. Enrichment of different BMSC ATAC-Seq peak types (stable, age-sensitive) over background among in different BMSC ChromHMM states is shown (Fisher’s exact test; * p-adjust < 0.1; **** p-adjust < 0.0001). Number of peaks indicated on the bars. **F)** Distributions of differential accessibility (log2 fold-change old vs young) are shown for peaks in the different BMSC ChromHMM states. For each BMSC ChromHMM state, peaks are grouped into age-sensitive and background.

On the RNA level, we found only 12 differentially expressed genes using DESeq2 at 10% FDR from 14,199 expressed genes (**Figure S2A**). To ensure this lack of differential signal was not a result of hidden batch effects, we used surrogate variance analysis (SVA) and remove unwanted variation (RUV) method, to remove hidden confounding factors (Risso et al., 2014), both of which found no differentially expressed genes.

To assess age-related changes of RNA isoforms in BMSCs, we performed differential exon usage analysis by DEXSeq (see Methods) using the BMSC RNA-Seq data, and identified 1,025 differentially used exons at 10% FDR and a minimum log2 fold change of 0.5. Amongst the differentially used exons, 473 (in 335 genes) and 552 (in 465 genes) are differentially excluded and included with increasing age, respectively (**Figure S2A**).

To quantify the effects of ageing on the BMSC proteome, we used the data from our previous study, where the Spearman correlation of protein abundance with the donors’ age was used as a measure of age-specific changes in the BMSC proteome (Hennrich et al., 2018). Here, the age-specific changes are represented as the relative increase or decrease in abundance (%) across all the ages between 20 and 60 (**Figure S2A**). In total, 760 proteins are significantly affected by ageing (Spearman p-value <0.05), of which 447 are ageing-up and 313 are ageing-down (**Figure S2A**).

TF binding sites in genome-wide accessible chromatin regions can be used to estimate the global TF activity, an approach implemented in the computational tool called diffTF (Berest et al., 2019). To estimate age-related changes in BMSC TF activity, we carried out diffTF analysis using the ATAC-Seq data and identified 350 from 678 sequence-specific TFs exhibiting differential activity upon ageing in BMSCs at 10% FDR (**Figure S2A, Table S8**). Interestingly, we identified IRF3 as an age-sensitive TF, which was previously described to be an important TF for supporting HSC expansion, regenerative and engraftment capacity (Nakahara et al., 2019), indicating the potential misregulation of the bone marrow niche environment during ageing.

Overall, we identified 2,152 age-sensitive genes in the BMSCs across all the molecular levels that are significantly affected by ageing on at least one level. The majority of age-sensitive genes (>90%) are affected by ageing only on one molecular level (**Figure 2A**). This lack of connection between promoter chromatin accessibility and protein abundance is consistent with the poor correlation between chromatin accessibility changes in promoters and protein abundance (r = 0.0036, p = > 0.75) (**Figure S2B**).

The results here show that BMSCs ageing hotspots affect distinctive subsets of chromatin regions, transcript isoforms, protein abundances and TF activities, but generally not the RNA levels. The lack of correlation between RNA levels and protein abundances could be explained by the post-transcriptional co-regulation of protein complex subunits. The complex subunit that is expressed (RNA level) at a higher amount than the other subunits, are often degraded (protein level) if they are not incorporated into the protein complex (Liu et al., 2017). The lack of correlation between chromatin signal and gene expression levels suggests an additional buffering mechanism between chromatin and transcription. This leads to the question of how ageing hotspots at each molecular layer affect each other and whether this contributes to changes in cellular function and potential.

### Post-transcriptional changes in ageing affect cell function

We characterised the functional impact of the 2,152 age-sensitive genes using Gene Ontology (GO) enrichment analysis and found distinct biological processes enriched among each molecular level. As observed previously (Hennrich et al., 2018), age-sensitive genes with downregulated protein abundances are enriched in chromatin organisation and RNA regulatory processes, while those upregulated are enriched in autophagy and extracellular membrane organisation processes (**Figure 2C**). Extracellular matrix organisation processes were found enriched for age-sensitive genes at the protein abundance and RNA isoform level, suggesting altered abundances or ratios of protein isoforms which could affect the bone marrow niche environment.

In agreement with the GO enrichment analysis on the protein level and RNA isoforms, age-sensitive proteins assigned to autophagy-related (Golgi vesicle transport & autophagy) and extracellular matrix organisation processes, are predominantly upregulated during ageing, while age-sensitive proteins that regulate cell cycle (centromere & cell cycle proteins), translation, alternative splicing (splice factors), polymerase II transcription, and chromatin remodelling (epigenetic factors) processes, are predominantly downregulated (**Figure S3D**). As a validation, we further studied some of the protein abundance changes in protein complexes (**Figure S3E**) that are relevant to the enriched GO processes. In general, the protein abundances in the complexes show very coherent up- or down-regulation (**Figure S3E**), suggesting that the entire molecular machine rather than individual subunits are age-sensitive.

Notably, 7 of the 13 GO biological processes enriched for age-sensitive genes on the protein abundance and RNA isoforms levels, are also enriched for genes that are significantly stable during ageing on the RNA and chromatin accessibility level (**Figure 2C, Table S1**, see Methods). Together with the observation that changes on the protein level correlates poorly with those on the chromatin and transcription levels (**Figure S2B**), the results here corroborate the notion that ageing affects protein abundance by affecting post-transcriptional regulation, independent of chromatin and transcriptional changes.

To further characterise the link between the changes in splicing factor abundance (**Figure S3D**) and differential exon usage upon ageing (**Figure S2A**), we studied the splice factor binding sites at close proximity (+ and - 200 bp respectively) to the 5’ and 3’ splice sites of the differentially used exons compared to that of the exons not differentially used in ageing. We found 72 RNA binding proteins whose binding sites are enriched in the close proximity of the differentially used exons at 5% FDR, of which 16 are age-sensitive proteins (**Figure S3F**). The downregulation of the splicing factors protein abundance in ageing could be responsible, at least in part, for the RNA isoform changes observed. Among the age-sensitive genes affected on the RNA isoform and protein abundance levels are ITGA5, COL4A2, PTK2 (FAK) and MAP4K4 (**Figure 2D**). In particular, ITGA5, COL4A2 and FAK together belong to the integrin signalling pathway (Hynes, 2002), strongly implicating their role in age related changes in the BMSCs.

### Misregulation of chromatin accessibility in ageing affects cell differentiation potential

While protein abundance and RNA isoform changes represent alterations in a cell’s functional state, chromatin accessibility changes as measured by ATAC-seq could alter the cell’s potential to respond upon stimulation or differentiation (Rendeiro et al., 2020). To investigate the BMSC differentiation potential, we stratified the ATAC-seq peaks into the ChromHMM chromatin states, which reflect genomic regions of distinct regulatory activity. We found age-sensitive chromatin regions strongly enriched for bivalent regions (OR=3.5), heterochromatin (OR=3.5), zinc-finger and repeat regions (OR=9.8), and moderately enriched in repressed polycomb (OR=1.8) and active transcription (OR=1.2) regions (**Figure 2E, S3A, S3C**). Bivalent/repressed polycomb domains regulate the process of stem cell differentiation by allowing the required gene expression programmes to rapidly respond to differentiation stimuli (Voigt et al., 2013). The enrichment of age-sensitive chromatin in bivalent and repressed polycomb regions, indicates ageing affects BMSC differentiation potential. In contrast, chromatin regions containing active transcription start sites are depleted from age-sensitive peaks (OR=0.66) and instead are strongly enriched for stable regions in ageing (**Figure 2E**; Methods). This suggests that the chromatin regions directly governing polymerase II transcription of RNA remains largely stable during ageing, and could represent the buffering mechanism that explains the lack of age-sensitive genes on the RNA level (**Figure S2A**) and the weak correlation between chromatin accessibility changes in promoters and RNA levels in BMSCs (r = 0.009, p = 0.24; **Figure S2B**). The ChromHMM states distribution of ATAC-Seq chromatin regions in BMSCs are robust with regards to changes in the consensus peaksets (**Figure S3B**).

The enrichment of age-sensitive regions in bivalent, heterochromatin, repressed polycomb, and zinc-finger genes and repeats regions, which are crucial for stem cell maintenance and function (Zhou et al., 2011), is predominantly associated with chromatin that closes upon ageing (**Figure 2E-F, S3A**), indicating a loss of stem-cell potential during ageing. Furthermore, genes with ageing-down promoter peaks are enriched for developmental processes (**Figure 2B**), indicative of an impaired stem cell differentiation potential in BMSCs with age.

Overall, ageing shows a two-pronged effect on the BMSC gene regulatory landscape. The biological processes affected by age-related protein abundance and RNA isoform changes (**Figure 2B**) in BMSCs represent well known hallmarks of ageing such as epigenetic misregulation and loss of proteostasis (López-Otín et al., 2013), recapitulating the universally known cellular ageing processes. Age-related chromatin changes (**Figure 2E**), on the contrary, are associated with the alterations of cellular differentiation potential in BMSCs. Amongst the age-sensitive changes (**Figure 2A**), 64% affect the BMSC post-transcriptional landscape including RNA isoform and protein abundance without affecting RNA expression levels and chromatin accessibility, whilst 31% affect the chromatin landscape without affecting gene expression components. Chromatin changes bear a “universal” pattern of affecting stem cell regulatory domains, a phenomenon also observed in other cell types (Liu et al., 2013; Rakyan et al., 2010), thereby changing the “potential” of the cell to answer to perturbation or challenges. We next investigate the effects of ageing on the “reactome” or “pertubome”, and hence ultimately the cell fate of BMSCs.

### Ageing-specific priming of early time point adipogenesis by age-sensitive TFs

We further investigated how ageing may alter the BMSCs’ differentiation into adipocytes and osteoblasts. Differentiation processes are typically driven by TFs, therefore we annotated age-sensitive TFs as adipogenesis or osteogenesis-related based on their GO biological processes annotation (Ashburner et al., 2000; The Gene Ontology Consortium, 2019) (**Table S2**). Of the 41 age-sensitive TFs, 16 are involved in adipocyte-, 20 in osteoblast-differentiation, and 5 in both (**Figure 3A**). Notably, while all 41 TFs were captured by RNA-seq, only 17 TFs are measurable by mass spectrometry (**Figure 3A**). We found adipogenesis-associated TFs show increasing activity with age, while osteogenesis-associated TFs show decreasing activity (**Figure 3B**).

**Figure 3.**
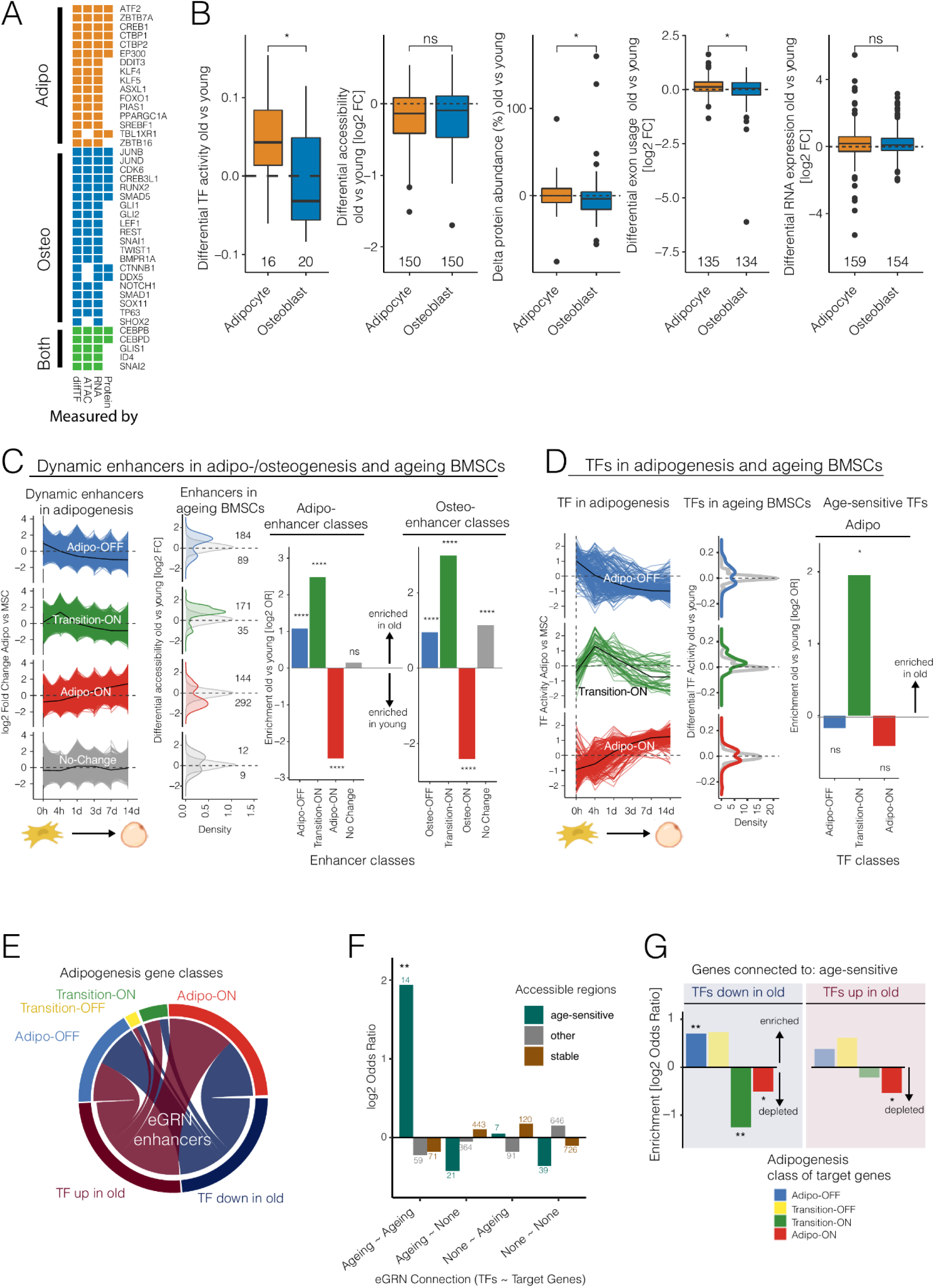
TF activity and chromatin accessibility at enhancers prime BMSC differentiation potential in ageing. **A)** The measures by which a TF is captured are shown for TFs annotated in GO terms related to adipogenesis (orange), osteogenesis (blue), or both (green). **B)** Differential signal between old vs young is shown for genes stratified based on whether they are annotated in GO terms related to adipogenesis (orange) or osteogenesis (blue) on the different molecular layers. From left to right: age-sensitive TFs (differential TF activity), promoter ATAC-Seq peaks in all genes (differential accessibility), protein abundance for all genes, alternative exon usage for all exons, and RNA expression level for all genes (Wilcox test, * p < 0.05). **C)** Left: chromatin accessibility of dynamic enhancers during adipogenesis differentiation time course taken from (Rauch et al., 2019) is shown as log2 fold change vs undifferentiated BMSCs at each time point. Each line represents one dynamic enhancer. Enhancers are split into four groups based on their pattern during differentiation: Adipo-OFF, Transition-ON, Adipo-ON, No Change. Middle: distribution of differential accessibility in old vs young BMSCs from our data is shown as log2 fold change and stratified by the enhancer class of dynamic enhancer that overlap with the respective peaks. Only ATAC-Seq peaks whose genomic positions overlap with dynamic enhancers are shown. Age-sensitive peaks are bold colors, non-ageing peaks in light grey. Right: enrichment of ATAC-Seq peaks up vs down in ageing BMSCs that overlap with enhancers in adipo- and osteogenic differentiation (Fisher’s exact test; n.s. not significant; **** p-adjust < 0.0001; Figure S5). **D)** Left: TF activity changes in adipogenic differentiation based on adipogenesis time course data from (Rauch et al., 2019). TFs are grouped into three classes based on their dynamic during differentiation: Adipo-OFF, Transition-ON, Adipo-ON. Middle: distribution of differential TF activity during ageing (age-sensitive TFs in colour, other TFs in grey). Right: enrichment of age-sensitive TFs up vs down in ageing BMSCs in TF classes. **E)** eGRN obtained from chromatin accessibility and RNA-seq data of BMSC adipogenic differentiation. Connections (eGRN enhancers) depict the relation between age-sensitive TFs and their target genes. The clusters of target genes are grouped into four classes: Adipo-OFF, Transition-OFF, Transition-ON and Adipo-On, according to their gene expression level changes during the course of the differentiation (Figure S7). **F)** Enrichment of age-sensitive accessible regions among eGRN enhancers stratified by age-sensitivity of the TF and target gene they connect. Colours indicate the age-sensitivity state of accessible regions (Fisher’s exact test, ** p-adjust < 0.01). **G)** Enrichment of genes connected to age-sensitive TFs that are up (left) or down (right) in ageing among the different gene classes from E (Fisher’s exact test, * p-adjust < 0.1, ** p-adjust < 0.01).

Among the age-sensitive TFs we observe well known regulators of the BMSC adipo- and osteo-differentiation, including PPARGamma, RUNX2, CEBPs, NOTCH1, CREB and CTNNB1 (Huang et al., 2015). A similar analysis for adipo- and osteogenesis related genes on the other molecular layers revealed no significant changes for age-sensitive genes, and only minor changes in protein abundance and differentially used exons when considering all genes regardless of whether they are age-sensitive or not (**Figure 3B & S4F**).

The fact that the increase in TF activity during adipogenesis does not lead to significant changes on RNA and protein levels, are consistent with the view that the BMSCs are still undifferentiated. However, they seem to be already lineage-primed on the TF activity and global chromatin levels, suggesting the cells will more readily differentiate into adipocytes over osteoblasts in the old age.

### Gene regulatory network analysis identifies global mechanisms of age-dependent BMSC differentiation bias

To further investigate the mechanisms of BMSCs lineage-priming during ageing, we integrated our data with a recently published BMSC adipo-osteo differentiation time course dataset (Rauch et al., 2019). The differentiation time course data consists of chromatin accessibility (DNase hypersensitive sites), RNA expression, and TF activity measurements at 6 time points (0h, 4h, 1d, 3d, 7d and 14d) during adipogenesis or osteogenesis (Madsen et al., 2018; Rauch et al., 2019). We re-processed the chromatin accessibility data to match our analysis pipeline, and defined *dynamic enhancers* involved in BMSC differentiation according to the authors’ definition (Rauch et al., 2019). We replicated the findings of the original study that in general more extensive remodeling is required for adipogenesis on the chromatin and RNA level compared to osteogenesis (Rauch et al., 2019) (**Figure S4A-E**).

We grouped the dynamic enhancers and TFs from the differentiation time course according to their signal (for enhancers)/activity (for TFs) across the differentiation into several classes using partitioning around medoids followed by visual inspection: “Adipo-OFF” (signal/activity decreases during adipogenesis), “Transition-ON” (signal/activity first increases and then decreases during adipogenesis), “Adipo-ON” (signal/activity increases during adipogenesis), and for enhancers also “No Change” (no change in signal/activity during adipogenesis) enhancers and TFs (**Figure 3C-D**). Similar classifications were done for the osteogenesis differentiation time course (**Figure S5-S6**).

To understand how changes in BMSCs ageing may impact their differentiation potential, we overlapped the dynamic adipogenic enhancers with our ageing ATAC-seq data. We found that ageing-up chromatin regions are strongly enriched in the “Transition-ON” enhancer class for adipogenesis, suggesting a role of chromatin priming in early adipogenesis events (**Figure 3C**). Since enhancers in the Adipo-ON (and similarly Osteo-ON) enhancer class are unlikely active in the undifferentiated BMSCs in this study, the enrichment of enhancers in the “Adipo-ON” enhancer class in young individuals corroborate the earlier observation of the loss of bivalent region accessibility in BMSCs during ageing (**Figure 3C**). A similar analysis for osteogenic differentiation (**Figure S5)** revealed also depletion of ageing-up regions in the “Osteo-ON” enhancer class. Examination of TFs in adipogenesis revealed strong enrichments for ageing-up TFs among the “Transition-ON” class (**Figure 3D)** while no significant association was found for age-sensitive TFs in any TF class obtained from osteogenic differentiation pattern (**Figure S6**). The association of age-sensitive chromatin and TFs with early adipogenesis events suggest a general loss of early time point differentiation potential of aged BMSC, which is in line with the closing of bivalent chromatin regions with age observed above

Since we observed an adipocyte early differentiation skew for both TFs and enhancers but no differentially expressed genes in our BMSCs, we hypothesised that the altered activity of the TFs and their impact on chromatin may prime BMSCs from older individuals towards adipogenic differentiation, while gene expression still reflects the current cellular identity.

To test this hypothesis, we constructed an enhancer-mediated gene regulatory network (eGRN) that allows us to characterize the target genes of age-sensitive TFs in terms of their expression dynamic during adipogenesis. Briefly, we applied our previously published algorithm (Reyes-Palomares et al., 2020) to the chromatin accessibility and gene expression data from the BMSC adipo-osteo differentiation time course data (Rauch et al., 2019), drawing TF-enhancer and enhancer-gene links based on co-variation across time points and replicates (see Methods). Our final eGRN comprises 160 TFs and 2034 target genes, forming 3066 TF-gene connections mediated by 1269 accessible chromatin regions (eGRN enhancers) at 20% FDR (**Figure 3E**). TF-gene pairs in which both of TF and gene are age-sensitive were strongly enriched for being connected by an age-sensitive ATAC-seq peak (**Figure 3F**) indicating age-sensitive TFs act as a mediator to propagate the age-sensitive chromatin signal through enhancers to its target gene.

Following the age-dependent adipogenesis-priming hypothesis, we speculate that ageing TFs would be connected to genes important in adipogenesis. To assess the hypothesis, we classified the genes based on their expression during adipocyte differentiation into “Adipo-OFF”, “Adipo-ON”, “Transition-ON” and “Transition-OFF” (expression first decreases and then increases during adipogenesis) classes similar to the dynamic enhancers and TFs described above (**Figure S5-S7**), and found that TFs more active in younger individuals are enriched for being connected to genes in the “Adipo-OFF” class and depleted from the “Adipo-ON” class (**Figure 3G**). The results indicate that a loss of activity in TFs connected with “Adipo-OFF” genes with age could lead to the downregulation of genes programmed to be shut off during adipogenesis, thus accelerating the differentiation process, in agreement with the initial hypothesis that TFs in older individuals could be primed towards adipogenesis. Examples of these TFs include known adipogenic TFs SNAI2 and CREB1 (Rauch et al., 2019).

Altogether, the eGRN revealed that age-sensitive chromatin regions could lead to the physiological differentiation bias in BMSCs differentiation by controlling the genes that are programmed to be shut-off in adipogenesis.

### Age-sensitive TFs and chromatin regions in BMSCs are enriched for SNPs associated with immune-related diseases/traits

Ageing is a major risk factor for many common diseases. To investigate how BMSC-ageing may contribute to complex traits and diseases, we integrated the age-sensitive genes with genetic variants that have previously been associated with common traits and diseases through genome-wide association studies (GWAS). Using the stratified linkage disequilibrium score regression (LDSC) enrichment method (Bulik-Sullivan et al., 2015) (see Methods) we identified 26 complex diseases/traits enriched among age-sensitive genes. The most enrichment we observed in age-sensitive TFs (**Figure 4A**), in line with recent observations that trans-regulators may be key drivers of diseases-associated processes (Freimer et al., 2021; Võsa et al., 2018). In addition to the expected BMSC-related traits, such as body height, bone density, and body mass index (BMI) traits, we identified several immune system-related traits and diseases, including asthma, allergy, inflammatory bowel disease, Crohn’s disease, and white blood cell measurements.

**Figure 4.**
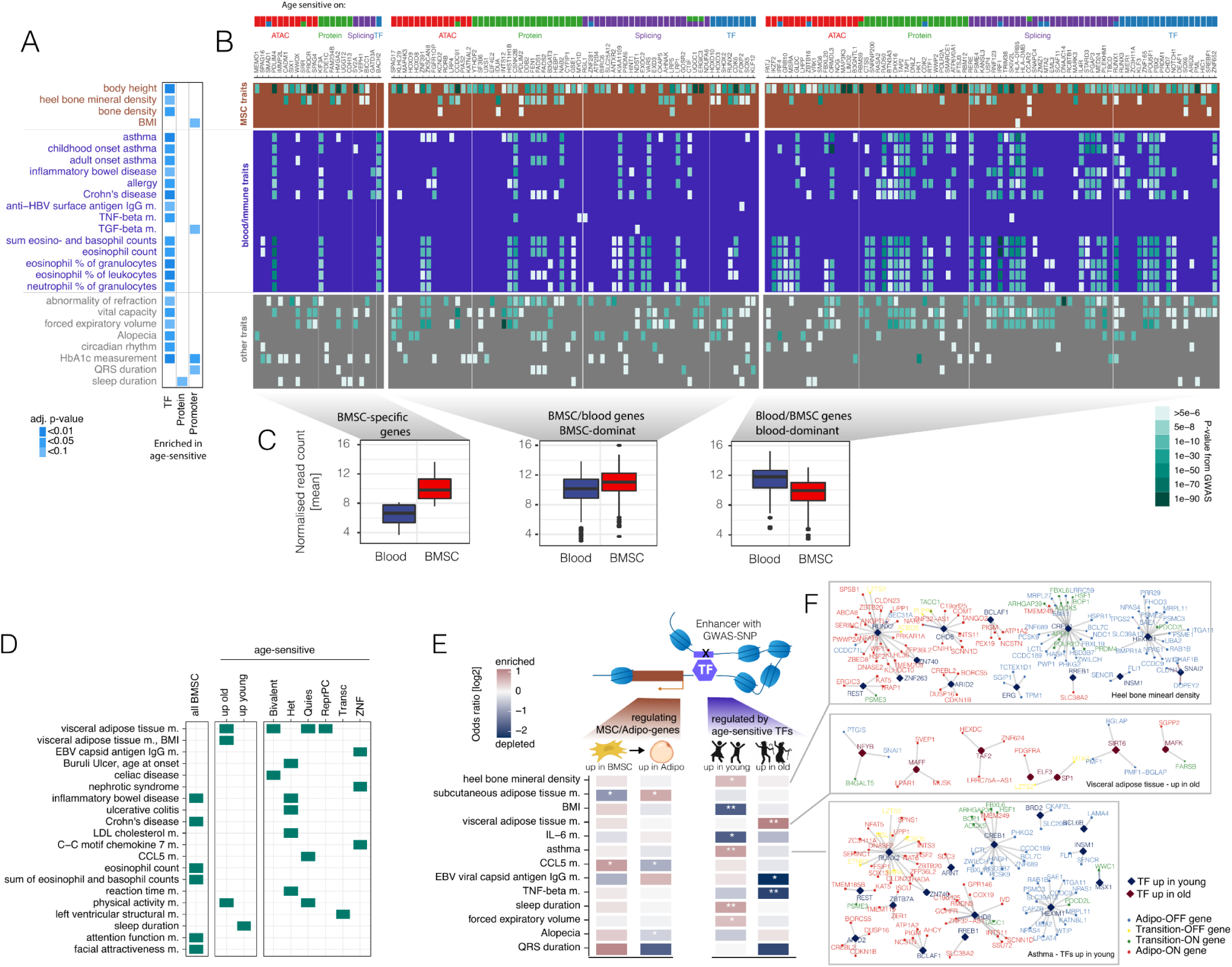
BMSCs ageing and genome-wide association studies traits/diseases. **A)** The heritability enrichment of trait/disease-associated SNPs among age-sensitive genes is shown. Trait-linked SNPs are obtained from the GWAS catalogue, genes are stratified by molecular level at which they are age-sensitive. SNPs are annotated to a gene if they lie within +/- 10kb of the gene body. **B)** Overlap of age-sensitive genes with SNPs (+/- 10kb) from the enriched traits in A are shown as the raw p-value per SNP in each GWAS. Traits/diseases are divided into 3 main categories, MSC-related traits (brown), blood/immune traits (blue) and other traits (grey). Age-sensitive gene categories are shown on top. **C)** Age-sensitive gene expression levels of human blood cells (from GTEx) and BMSCs. Genes are categorised into BMSC-specific, BMSC-dominant and blood-dominant genes based on gene expression levels in the two tissues. **D)** The heritability enrichment of traits/diseases-associated SNPs is shown for different age-sensitive ATAC-Seq peak categories (green boxes indicate p-adjust < 0.1). **E)** Left: enrichment of trait associated SNPs in eGRN enhancers linked to genes that change expression during adipogenic differentiation (up in BMSC=Adipo-OFF class; up in Adipo=Adipo-ON; classification from SFigure 7) (Fisher’s exact test, * p-adjust < 0.1). Right: enrichment of trait-associated SNPs in eGRN enhancers regulated by age-sensitive TF (Fisher’s exact test, * p-adjust < 0.1, ** p-adjust < 0.01). **F)** Examples of eGRN in selected traits. Diamonds represent TFs, small squares represent target genes. Colours indicate age-sensitive TF activity, and adipogenic differentiation category.

To further investigate the cell type specificity of the enriched immune traits, we selected age-sensitive genes in a +/- 10kb region from the top 15 GWAS SNPs with the lowest p-values per enriched trait (**Figure 4B**) and divided the genes into three groups based on their expression level in BMSCs and whole blood from GTEx data: (i) *BMSC-specific genes* - very high expression in BMSCs and very low expression in blood, (ii) *BMSC-dominant genes* - higher expression in BMSCs than in blood, and (ii) *blood-dominant genes* - higher expression in blood than in BMSCs (**Figure 4C**). Notably, all genes are highly expressed in BMSCs across donors.

While the blood-dominant genes associate with more GWAS SNP regions in immune traits, a number of immune trait-associated genes were either BMSC-specific or BMSC-dominant, indicating that the ageing of BMSCs may contribute to altered immune system function. Several age-sensitive genes are associated with both bone- and immune-related traits, potentially reflecting BMSC-intrinsic processes underlying their immune-modulatory or HSPCs support function. One example is PDLIM4, a bone- and immune-trait associated age-sensitive gene specifically expressed in BMSCs, that plays important roles in bone development and osteoporosis (Charoenpanich et al., 2014). Its link to many immune related processes through age-sensitive ATAC-seq regions suggests that BMSC-intrinsic processes could regulate immune system function in the bone marrow niche environment. Another interesting example is HMGA2, known for its role in driving adipogenesis (Xi et al., 2016), which is associated with bone mineral density and asthma.

A similar enrichment analysis among the age-sensitive chromatin regions, stratified into chromatin states, revealed 20 traits, including several BMSC-related physical traits, such as “visceral adipose tissue”, body mass index (BMI) and BMI-correlated traits such as “physical activity” and “sleep measurement” (Jones et al., 2016; Klimentidis et al., 2018) (**Figure 4D**). BMSC chromatin regions not associated with ageing are enriched for immune-function related diseases/traits, again highlighting the potential contribution of BMSCs to immune system regulation in the bone marrow niche. Age-sensitive bivalent, quiescent and repressed polycomb regions are enriched for “visceral adipose tissue”, further corroborating the effects of ageing on stem cell function and adipogenesis. Bivalent and heterochromatin are enriched for immune-related, metabolic and physical traits and diseases, including “low density lipoprotein cholesterol”, “reaction time (cognitive) measurement”, “inflammatory bowel disease” and “ulcerative colitis”. Zinc-finger and quiescent regions are further enriched for blood and immune traits.

The associations of BMSC age-sensitive chromatin regions with various immune phenotypes, strongly support the role of BMSCs in age-related immune changes in the bone marrow niche environment.

### Down-regulation of age-sensitive TFs in BMSCs could orchestrate systemic immune traits

To understand the wider impact of BMSC adipogenesis and the age-dependent effects on chromatin and TF priming in the context of adipogenesis (**Figure 3**), we next investigate i) how eGRN enhancers in adipogenesis could contribute to common traits and diseases, and ii) how age-sensitive TFs in the context of the eGRN associate with common traits and diseases. We first overlapped the eGRN enhancers with GWAS SNPs (p-value < 5e-03) of the traits that we found enriched among age-sensitive genes and peaks from **Figure 4A**,**D**. For each trait, we then quantified whether its associated SNPs are enriched in i) eGRN enhancers connected to Adipo-ON or Adipo-OFF genes (**Figure 3E left**), indicating the contribution of the BMSC differentiation program, or ii) in eGRN enhancers regulated by age-sensitive TFs, implicating age-dependent priming of BMSCs, to the corresponding common traits (**Figure 3E right**).

We found trait-associated SNPs enriched among enhancers of the BMSC differentiation programme for two traits: enhancers of “Adipo-ON” genes (programmed to be turned on in adipogenesis) for subcutaneous adipose tissue measurement, and enhancers of “Adipo-OFF” genes (programmed to be turned off in adipogenesis) for CCL5 measurement. Among eGRN enhancers regulated by age-dependent priming we found trait-associated SNPs of five traits enriched, including traits related to BMSC-derived cells. For example, ‘heel bone mineral density”-associated SNPs are enriched for being regulated by TFs with activity higher in young individuals, whereas SNPs associated with “visceral adipose tissue” are enriched for being regulated by TFs with activity higher in old individuals, further corroborating our observations above that ageing primes adipogenic potential (**Figure 4E-F**). Interestingly, age-primed enhancers are also enriched for asthma, an immune-related trait, in the young, which is consistent with previous findings that found allergic asthma more frequently in younger individuals (Pakkasela et al., 2020).

BMSC transplantation as a therapeutic option in animal models of asthma strongly indicates the immunosuppressive function of BMSCs in alleviating asthamtic symptoms (Mirershadi et al., 2020). The results here (**Figure 4E**), together with our observation of age-sensitive genes and peaks enrichment in immune-related traits (**Figure 4A-D**), strongly indicate that age-dependent priming of BMSC differentiation could simultaneously alter the immune-regulatory pathways shared, contributing to altering the bone marrow niche environment. The eGRN allowed us to study how age-sensitive TFs may enhance the effect of common trait-associated SNPs during ageing and to gain insights on the genes they may regulate (**Figure 4F**).

Altogether the results here indicate that both differentiation-related cellular changes in BMSCs as well as alterations of BMSC function during ageing, could contribute to the changes in inflammatory conditions, various physical and metabolic changes on the systemic level via enhancers associated with trait- or disease-associated common polymorphisms.

## Discussion

Bone density loss during normal physiological ageing has long been known to associate with BMSC differentiation bias as a result of increased bone marrow adipocytes and decreased osteoblasts formation (Justesen et al., 2001; Meunier et al., 1971; Moerman et al., 2004). While several studies have investigated the effects of ageing on different signalling pathways that could contribute to the BMSC adipo-osteogenic differentiation bias (Almeida et al., 2009; Stolzing et al., 2008; Sun et al., 2006), the underlying mechanisms remained poorly understood. The results of our study suggest a molecular mechanism, by which age-sensitive TFs prime BMSCs towards early adipogenesis already in their MSC fate. Overall, our study on the ageing of bone marrow mesenchymal stromal cells (BMSC) has shed light on the molecular mechanisms underlying the physiological ageing of the bone marrow niche.

The multiomics angle of our study allowed us to investigate the impact of ageing in BMSCs on three different regulatory levels: chromatin, RNA, and protein. Amongst the different gene regulatory levels in our datasets, changes in protein abundance represent the most direct readout for variations in biological pathways activities, while chromatin accessibility and TF activity inferred from chromatin accessibility are more reflective of the differentiation potential of the cells. Previous studies on bone marrow niche ageing have focused on proteomics and RNA-seq and have identified processes such as extracellular matrix remodeling being affected during ageing (Hennrich et al., 2018). While TFs represent a readout of the differentiation potential of BMSCs, they are highly underrepresented in proteomics experiments, thus highlighting the importance of integrative analyses, such as profiling chromatin accessibility and TF activity (diffTF analysis) to characterise TF functions (Weidemüller et al. 2021).

On the chromatin level, bivalent/repressed polycomb domains regulate the process of stem cell differentiation by allowing the required gene expression programmes to rapidly respond to differentiation stimuli (Voigt et al., 2013), while zinc-finger genes often directly regulate the expression or act as co-factors of the TFs in differentiation (Cassandri et al., 2017). Heterochromatin mediates various biological processes including centromere function and cell differentiation (Allshire and Madhani, 2018). The destabilisation of heterochromatin is a major hallmark of ageing and cellular senescence, contributing to increasing genome instability and transposable element activity (Criscione et al., 2016; O’Sullivan and Karlseder, 2012). The enrichment of differentially accessible peaks in bivalent, heterochromatin and zinc-finger regions suggests that many age-specific chromatin accessibility changes in the BMSC specifically affect the regulatory elements for stem cell function during ageing.

The fact that most age-sensitive genes are only present on one molecular layer highlights the complexity of biological ageing. Our observations that very few genes are age-sensitive on the transcriptomic level, yet pervasive changes occur on the chromatin and protein level, could be explained by the enrichment of “stable” chromatin peaks observed on the majority of the TSS, and the general age-specific alterations in most transcriptional related processes on the proteome level (Liu et al., 2017). This indicates that the effects of ageing on transcription are unlikely to happen in a unified and predictable manner. In a previous study we found that differential RNA expression is predictable by variation of RNA levels across individuals (Sigalova et al., 2020). Thus, an alternative explanation for the lack of age-sensitive genes on the RNA-level could be that the heterogeneity of the ageing cohort masks whatever small signals are there. The overall downregulation of the proteome responsible for chromatin organisation, including components of the polycomb group complex that regulates the repression of bivalent chromatin regions, could explain the strong changes observed at the chromatin ageing-hotspots that are not translated to the RNA level which is presumably at steady state.

The bone marrow and BMSCs have been shown to directly affect the common traits enriched (**Figure 4**), including inflammatory conditions (Allakhverdi et al., 2013; Yagi et al., 2010), granulocyte traits (Brigger et al., 2020), bone density, and other physical traits. With the results of our study, we can now propose hypotheses about the mechanisms by which common polymorphisms in BMSCs may play a role in immune-regulatory pathways. Specifically, we can use our eGRN to identify the genes that may be targeted by these SNPs and the TFs that may modulate their activity.

The bone marrow is recognised to function on a systemic level as a lymphoid organ that has immune regulatory roles (Zhao et al., 2012). Immune cells from both the adaptive and innate immune system reside and migrate into and out of the bone marrow, where both BMSCs and various immune cells regulate each other and the bone marrow microenvironment via cell-cell contact and/or soluble factors (Zhao et al., 2012). MSC-derived osteoblastic cells can also regulate the function (Calvi et al., 2003), homing after transplantation (Lo Celso et al., 2009) and lineage maturation (Ding and Morrison, 2013; Yu and Scadden, 2016; Zhu et al., 2007) of HSPCs, while adipocytes reduce hematopoietic activity (Naveiras et al., 2009) and inhibit lymphoid differentiation (Bilwani and Knight, 2012). Thus, in addition to affecting bone density, the age-induced balance shift from osteocytes to adipocytes may also have a crucial role in HSC maintenance, evidenced by the associations of age-related MSC changes in the bone marrow niche environment with the misregulation of HSC function and differentiation, potentially contributing to undesirable haematological changes such as cancer development (Kovtonyuk et al., 2016).

GWAS have revealed hundreds of thousands of genetic variants associated with common traits and diseases. Ageing, and specifically the ageing of the immune-system are likely to contribute to many common diseases. Indeed, our study revealed several common traits and diseases, including diseases related to altered immune system function are enriched for age-sensitive genes. By integrating the data from our ageing cohort at the level of an enhancer-mediated gene regulatory network, with several common and age-dependent diseases, our study provides a stepping stone towards investigating the molecular links between common immune-related traits and the ageing of the bone marrow stem cell niche.

## Methods

### Isolation of BMSCs from bone marrow

NMSCs were obtained as previously described (Hennrich et al., 2018; Pellagatti et al., 2018). Briefly, healthy donors were punctured at the posterior iliac crest using a Yamshidi needle. Aspirations of 10ml were taken at 5-7 different levels. The procedure took place at the University Hospital Heidelberg and the study was approved by the Ethics Committee for Human Subjects at the University of Heidelberg, and written informed consent has been obtained for each participant. The distribution of donors is shown in Figure 1A. These aspirates were washed with phosphate-buffered saline (PBS, Sigma-Aldrich #D8537). The eluates were laid on FICOLL-Paque (15 mL, Biochrom), centrifuged (800g, 30 minutes) to separate mononuclear cells (MNCs). The MNC fraction was seeded in T75 cell culture flasks with Verfaillie medium (VM) at a density of ∼10^6^ cells/mL, and cultured in fibronectin coated flasks until adherent colonies formed (at least 4 days). The adherent colonies were defined as passage 0 and for expansion of the cells, freshly isolated or thawed BMSCs were culture-expanded in VM for one passage and then seeded in parallel into VM at 37 °C with 5% CO2.

### ATAC-seq library preparation

ATAC-Seq library was prepared by either the “Fast-ATAC” protocol (Corces et al., 2016) or the “low mitochondrial read” protocol (Litzenburger et al., 2017) as previously described. Fifty thousand BMSCs were harvested at passage 4 and were washed with 50 μL of ice cold PBS. Lysis and transposition were performed according to the protocol used.

For “Fast-ATAC”, lysis and transposition reactions were performed simultaneously in 50 μL buffer (25 μL 2x TD buffer, 2.5 μL Tn5 Transposase TDE1, 0.5 μL 1% digitonin, 22 μL nuclease-free-water) (Illumina FC-121-1030 Nextera DNA library prep kit). For “low mitochondrial read” protocol, cells were simultaneously lysed and nuclei pelleted in 50 μL cold lysis buffer (10 mM Tris-HCl, pH 7.4, 10 mM NaCl, 3 mM MgCl2 and 0.1% IGEPAL CA-630, 0.1% Tween-20) at 500 rcf for 10 mins at 4°C. Immediately after lysis, DNA transposition was performed in transposition buffer (25μl 2x TD buffer, 2.5 μL TDE1, 22.5μl H2O, and 0.5 μl 1% Tween-20, 22 μL nuclease-free water) at 37 °C for 30 min.

Libraries were prepared by PCR amplification and barcoding as previously described (Buenrostro et al., 2015) using Illumina Nextera i7 and i5 primers. The prepared libraries were quantified and sequenced paired-end 75 bp on an Illumina NextSeq 500 at EMBL Genomics Core Facility.

### Processing of ATAC-seq data

Libraries were processed with our in-house ATAC-Seq data processing Snakemake pipeline as previously described (Berest et al., 2019). FastQC (https://www.bioinformatics.babraham.ac.uk/projects/fastqc/) was used to assess sequence quality. Adapter sequences were removed using Trimmomatic (Bolger et al., 2014) with the parameters ILLUMINACLIP:NexteraPE-PE.fa:1:30:4:1:true TRAILING:3 MINLEN:10. Sequencing reads were aligned to hg38 genome assembly using Bowtie2 (Langmead and Salzberg, 2012) with parameters --very-sensitive -X 2000, followed by various post alignment cleaning steps (Picard tools CleanSam, FixMateInformation, AddOrReplaceReadGroups, and ReorderSam) (https://broadinstitute.github.io/picard/).

Base quality recalibration was performed using GATK (McKenna et al., 2010) with known variants from the GATK bundle for hg38 (SNPs: dbSNP version 138, Indels: Mills_and_1000G_gold_standard.indels) to increase the data quality by correcting for systematic errors by the sequencer when estimating the quality score of each base call.

Further cleaning steps were performed, including mitochondrial reads removal, non-assembled contigs or alternative haplotypes removal, reads filtering with a minimum mapping quality of 10, removal of duplicate reads by Picard tools, start sites adjustment as previously described (Buenrostro et al., 2013), and insertions or deletions removal Samtools.

Peak calling was performed using MACS2 (Zhang et al., 2008) (--no lambda, --nomodel, -qval 0.1, --slocal 10000) and blacklisted regions removed. Transcription start site (TSS) enrichment score and the irreproducibility discovery rate (IDR) (Li et al., 2011) were used to exclude samples with ATAC-Seq peaks of relatively poor reproducibility.

### Defining consensus peak set for ATAC-seq

Consensus peak set was generated from a total of 404,421 peaks using DiffBind (Ross-Innes et al., 2012). There are 36k (0.9%) shared across all 17 individuals and 34 replicate samples. As a final consensus peakset, the peaks were required to be present in at least 4 of the 34 samples, equivalent to 2 individuals, resulting in 140,635 peaks.

### ChromHMM chromatin state annotation

ATAC-Seq peaks were overlapped with mesenchymal stem cell DNase hypersensitivity data in the chromatin 15-state model annotation (E026_15_coreMarks_hg38lift_mnemonics.bed) and from ENCODE (Roadmap Epigenomics Consortium et al., 2015). The 15 states were further grouped into several general states for some analysis: Bivalent (TssBiv, BivFlnk, EnhBiv), Quiescent (Quies), Transcribed (TxWk, Tx, TxFlnk), Enhancer (Enh, EnhG), TssA (TssA, TssAFlnk), Heterochromatin (Het), ZNF_Rpts (ZNF/Rpts), and Repressed PC (ReprPCWk, ReprPC).

### Differentially accessible peak analysis

DiffBind (Ross-Innes et al., 2012) and DESeq2 (Love et al., 2014) bioconductor packages were used to calculate differentially accessible peaks. Peak-summits were recentered and extended +/-250bp. Different sequencing runs (i.e. Illumina flowcell) and donor genders were considered batch effects, and were removed in DESeq2 analysis. Differential analysis on the 140,635 consensus from a minimum overlap of 4 samples (Figure S1A) were calculated using an age-group study design (young: ≤ 29 years, middle: ≥30 ≤ 50, and old ≥51) in DESeq2 with the design (design = ∼Batch_effect + Age_group), parameters (test = “LRT”, reduced = ∼Batch_effect), following the standard workflow.

### Robustness analysis of differential ATAC-seq enrichment with chromHMM

The peak distribution in various chromatin states was found to be conserved when the differential peaks are identified from different consensus peaksets derived from the sequential removal of each individual in the differential analysis **(Figure S3B**), indicating that the differentially accessible peaks identified are robust.

### Processing of RNA-seq

RNA-seq data was obtained from (Hennrich et al., 2018; Pellagatti et al., 2018) and re-processed using DESeq2 using the standard workflow with the age-group study design (young: ≤ 29 years, middle: ≥30 ≤ 50, and old ≥51). The normalised RNA-Seq read counts per donor (Figure S1D), and the number of proteins characterised by mass spectrometry across donors (Figure S1E) were used to assess data quality. The variation of the normalised RNA-Seq read counts are comparable between donors, where the log10 median read counts of the samples centre around two (Figure S1D). SVA package (Leek et al., 2012) was used to further confirm the lack of signal following the standard workflow.

### Differentially used exon analysis

The transcript information from 29509 filtered genes was used for exon mapping using DEXSeq (Anders et al., 2012). To improve the biological relevance of the analysis, we applied a log2 fold change threshold of 0.5 for the differential exons, corresponding to a minimum 1.4-fold change, a deviation level higher than the 1.2-fold that is generally regarded as biologically relevant (Bray et al., 2003; Caffrey et al., 2008; Yan et al., 2002).

### Stable ATAC-seq /RNA-seq analysis

To complement the differential analysis, a stability analysis was performed to identify chromatin regions and RNA transcripts that are significantly more stable between old and young using DESeq2 with the following parameters: (lfcThreshold = 0.7, contrast = c(“Group”, “Young”, “Old”, altHypothesis = “lessAbs”). We identified 26,450 stable chromatin regions at 10% FDR and with a log2 fold change threshold of less than 0.7 (**Figure S2B**), while the expression of 710 genes that are stable at 10% FDR **(Figure S2B**). In addition to the stably expressed genes and stable chromatin regions, we found 8,323 genes to contain at least one stable ATAC-Seq peak in the promoter region, of which 134 genes also contain a differentially accessible promoter peak.

### diffTF analysis

TF binding sites used in this study are obtained from the HOCOMOCO and Remap databases (Chèneby et al., 2018; Kulakovskiy et al., 2013). TF activity was calculated using diffTF (Berest et al., 2019) using the following parameters (regionExtension 100, conditionComparison “Old,Young”, nPermutations 1000, nBootstrapts 0, nCGBins 10, minOverlap 4, regGenome Homo_sapiens_assembly38). Non-expressed TFs in BMSCs were removed from further analysis. We used 1% false discovery rate as a threshold for statistical significance. Different sequencing runs (i.e. Illumina flowcell) and donor genders were considered batch effects, and were removed in the analysis

### QC of proteins

Proteomics data were obtained from (Hennrich et al., 2018). A total of 6287 proteins were quantified by mass spectrometry. Briefly, proteins were quantified by tandem mass tag labelling, followed by liquid chromatography-tandem mass spectrometry analysis (Hennrich et al., 2018). Over 85% and 65% of the proteins characterised by mass spectrometry are present in more than 10 and 40 out of the 45 samples, respectively, indicating that the majority of the proteins are consistently and confidently identified across samples.

### Mapping of genes to reactome pathways

Reactome pathways (Fabregat et al., 2018) were grouped at the highest hierarchy, as previously described in (Hennrich et al., 2018). We then also added TF genes defined as the genes that have a specific DNA binding motif in the HOCOMOCO database (Kulakovskiy et al., 2013) (**Figure 1G**).

### GO term enrichment

Due to the similarities of many GO terms, both in their associated genes and semantics, similar GO terms were grouped together using clusterProfiler (Yu et al., 2012) and the most significant GO terms are displayed (**Table S1; Figure 2E**).

### GO IDs for differentiation

We defined genes involved in adipogenesis using the following GO ids: 0045599, 0045600, 0045598, 0050873, 0045444, 0090335, 0090336, 0050872, 1903444. We defined genes involved in osteoblast differentiation using the following GO ids: 0045669, 0045668, 1905241, 0001649, 0045667, 0002076, 0002051, 004339, 1905240.

### Splice factor and protein complex analysis

Splice factor motifs data were downloaded from MotifMap (http://motifmap.ics.uci.edu/) (Daily et al., 2011) (Xie et al., 2009). Motifs overlapping a 200bp 3’ and 5’ regions in introns were included in enrichment calculation. Enrichment was calculated by Fisher’s exact test, 5% false discovery rate for statistical significance. Protein complexes data were downloaded from CORUM database (https://mips.helmholtz-muenchen.de/corum/) (Giurgiu et al., 2019), only complexes with at least 10 proteins are included.

### Reprocessing of the RNA, DNase hypersensitivity, and TF activity data from Rauch et al

Data was obtained from (Rauch et al., 2019) (direct communication). We selected all differentially accessible DNase hypersensitive sites across all time points, and clustered their normalised counts using partitioning around medoids clustering into 12 groups. Similarly, we clustered the normalised read counts for the RNA data. For TF activity, we clustered the activity levels of the TFs provided by the authors. The resulting clusters were manually classified into “Adipo-OFF” if the accessibility was decreasing throughout the timepoints; “Transition-ON” if there was an early upregulation followed by a decrease; “Adipo-ON” if there was a consistent increase, “Transition-OFF” if there was an early decrease followed by an increase, and “No Change” if there was no obvious changes across the time points (**Figure S5-S7**). We followed the same approach for classifying enhancers, genes and TFs according to their behaviour during osteogenesis (**Figure S5-S7**)

### Overlapping BMSC and Rauch et al datasets

To overlap our ATAC-Seq peaks with the DHS peaks from Rauch et al, GenomicRanges was used and with a minimum overlap of 1bp. There are 44,533 overlaps (32% of ATAC-Seq peaks) in total. To overlap TFs in BMSCs with those in Rauch et al. data, ENSEMBL ids of the TFs were used for matching. There are 305 TFs (34% of diffTF results) matching in Rauch et al. data.

### Construction of the gene regulatory network (GRN)

Gene regulatory network was constructed as previously described (Reyes-Palomares et al., 2020). Briefly, DNase hypersensitivity and RNA data from Rauch et al. were used for linking enhancers to putative target genes by correlating the accessibility of enhancers with the expression of the putative target genes that are within 250 kb of the peak. Links between TFs and enhancers are identified by correlating TF expression and enhancer accessibility signal at peaks that contain the TF motif. Each enhancer therefore is linked to at least 1 TF and 1 target gene, establishing the TF to target gene relation. We used a 20% false discovery rate to remove results that are considered not statistically significant. Both TF motifs from the HOCOMOCO and Remap database were used.

### GWAS enrichment analysis by LDSC

We first obtained and reformatted a total of 853 GWAS datasets from the GWAS catalog (Buniello et al., 2019) by converting the summary statistics columns into a format required by the LDSC tool (https://github.com/bulik/ldsc) for enrichment analysis (Bulik-Sullivan et al., 2015; Finucane et al., 2015). Input peaks and gene set files were converted as required by the LDSC tool. p-value adjustment was performed individually for each trait/disease.

### GWAS-gene analysis

To ensure that the enrichment in immune traits is indeed specific to BMSCs and not dominated by genes mostly expressed in immune cells, we determined the tissue specificity of GWAS gene sets by comparing gene expression of our BMSCs with blood and adipose tissue from GTEx.

### P-value adjustment

Unless otherwise stated, all p-values adjustment by Bejanmin-Hochberg method.

### Ethics statement

The study has been approved by the Ethics Committee for Human Subjects at the University of Heidelberg, and written informed consent was obtained from each individual.

## Acknowledgment

This study is mainly financed by financed by Baden-Wuerttemberg Stiftung GmbH and supported by German Cancer Aid (grant number 70114435). We thank the EMBL Genomics Core Facility for their expert help in Next-Gen Sequencing. We thank Alexander Rauch for providing the differentiation BMSC datasets and support through personal communication. We thank all the European Molecular Biology Laboratory and Molecular Medicine Partnership Unit Groups for their discussion and support. The Genotype-Tissue Expression (GTEx) Project was supported by the Common Fund of the Office of the Director of the National Institutes of Health, and by NCI, NHGRI, NHLBI, NIDA, NIMH, and NINDS. The data used for the analyses described in this manuscript were obtained from: the GTEx Portal on 26/08/2019. The accession ID (GCST) of the GWAS Catalog studies are in **Table S2** downloaded from the GWAS Catalog on 28/09/2020.

## Supplementary Tables

Column names and the containing information are as follows in the format “column_name: containing_information”.

**Supplementary Table 1** contains information of all of the GO term enrichment used in Figure 3. ID: GO ID; Description: GO ID description; p.adjust; geneID: ENSEMBL ID for foreground genes in the GO term; allGenes: ENSEMBL ID for foreground genes in the GO term and all other GO terms that were grouped together based on similarity; type: gene omics type.

**Supplementary Table 2** contains eGRN connections with their enhancers overlapping with GWAS SNPs. from: TF name; to: target gene name; from.2: TF ENSEMBL ID; to.2: target gene ENSEMBL ID; ATAC_peak_id; RSID: GWAS SNP IDs; gwas_P: p values of SNPs in the GWAS; gwas_lowestP: lowest p value of SNPs; gwas_lowestP_id: RSID of SNP with lowest p value; gwasID: GCST ID from GWAS Catalog; peakType: ageing peak type of enhancer in eGRN: traitName: name of GWAS; traitType: type of trait classified in Figure 4.

**Supplementary Table 3** contains all age-sensitive genes and the omics level they belong. ATAC: 1 (age-sensitive) and 0 (not age-sensitive) on promoter ATAC-Seq level; Splicing: 1 (age-sensitive) and 0 (not age-sensitive) on RNA splicing level; RNA: 1 (age-sensitive) and 0 (not age-sensitive) on RNA level; Protein: 1 (age-sensitive) and 0 (not age-sensitive) on protein level; diffTF: 1 (age-sensitive) and 0 (not age-sensitive) on TF activity level; ensg: ENSEMBL ID; hgnc_symbol: gene symbol; count: on how many omics levels the gene is age-sensitive; type: which omics levels the gene is age-sensitive.

**Supplementary Table 4** DESeq2 results for chromatin accessibility data. log2FoldChange_old/young: log2 fold change of read counts in old vs young individuals; pvalue: DESeq2 p values; padj: p adjusted values; samplePeakID: ATAC-Seq peak coordinates as IDs; msc_peakType: age-sensitive as ageing, not age-sensitive as background.

**Supplementary Table 5** DESeq2 results for RNA-Seq data. log2FoldChange_old/young: log2 fold change of read counts in old vs young individuals; pvalue: DESeq2 p values; padj: p adjusted values; ensembl.id: ENSEMBL ID.

**Supplementary Table 6** DEXSeq results for differential exon usage. Ensembl_gene_id: ENSEMBL ID; featureID: exon ID; pvalue: p values from DEXSeq; padj: p adjusted values; log2fold_Old_Young: log2 fold change of read counts in old vs young individuals; transcripts: ENSEMBL transcript ID; type: age-sensitive as ageing, not age-sensitive as background.

**Supplementary Table 7** Proteomiccs data. Ensembl_gene_id: ENSEMBL ID; approv.sym: gene symbol; spearman.pvalue: spearman correlation p values; spearman.cor: spearman correlation values; relativeChange20To60(%): relative changes in protein abundance from 20 to 60 years old.

**Supplementary Table 8** diffTF results. Ensembl_gene_id: ENSEMBL ID; Transcription.factor: transcription factor names; TF_activity_old/young: TF activity old vs young; pvalueAdj: p adjusted values; pvalue: p values; database: TF motif database.

