## Supplementary Figures for "Enhancer-priming in ageing human bone marrow mesenchymal stromal cells contributes to immune traits"

S1A

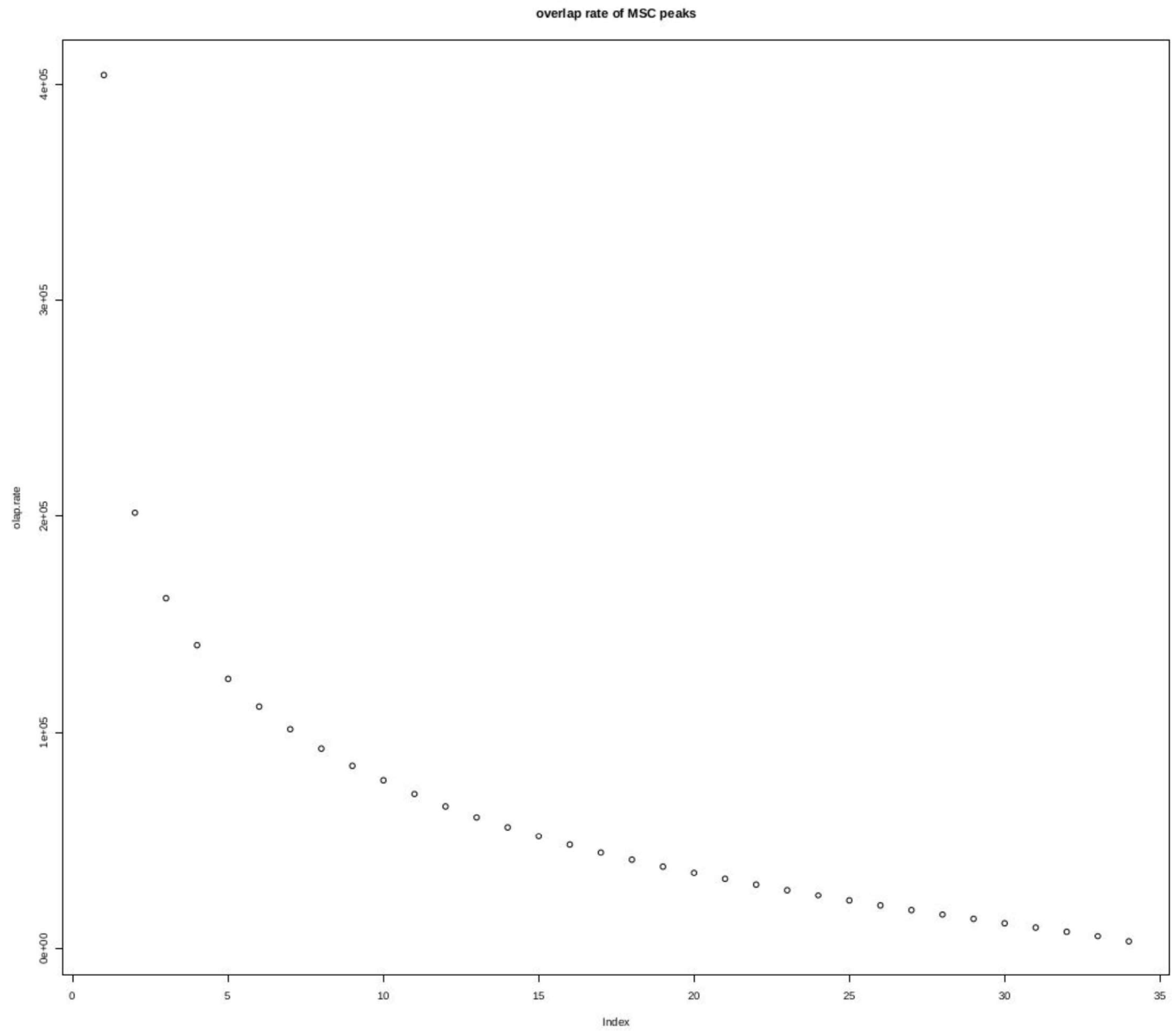


Figure S1A. Proportion of ATAC-Seq peaks overlaps in samples

S1B

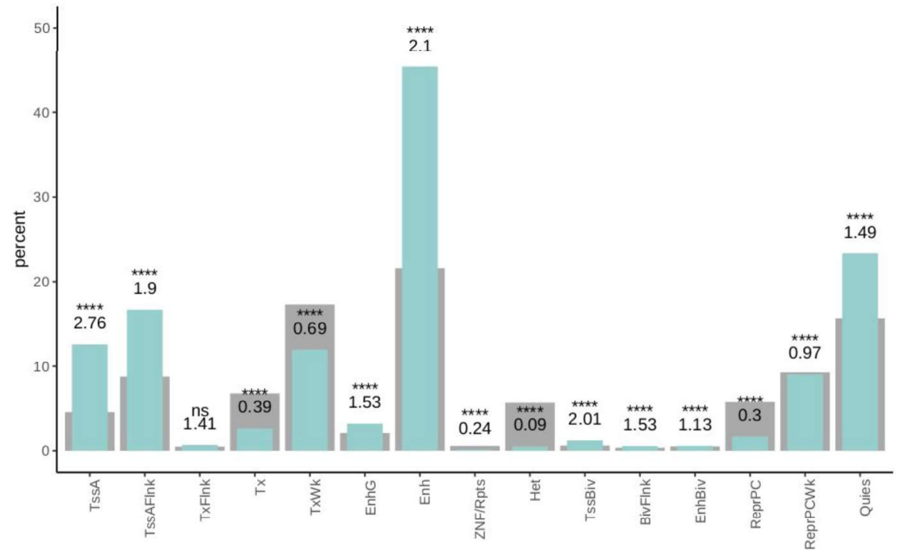


S1C

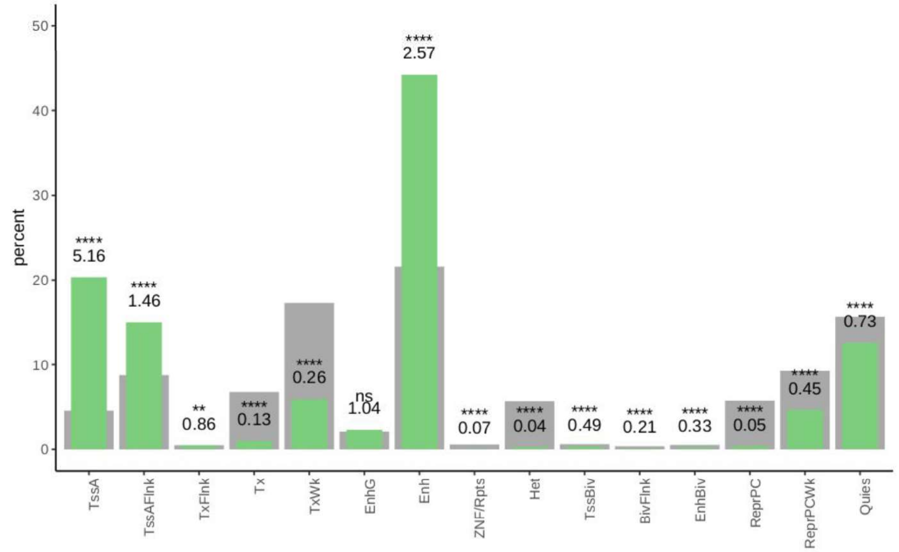


Figure S1B. Percentage of ATAC-Seq peaks in BMSC (blue), compared to chromHMM regions of mesenchymal stem cells (grey). Numbers indicate odds ratio (Fisher's exact test, **** p-adjust < 0.0001, ns not significant)

Figure S1C. Percentage of DNase hypersensitive sites in of human mesenchymal stromal cell data from ENCODE (green), compared to chromHMM regions of mesenchymal stem cells (grey). Numbers indicate odds ratio (Fisher's exact test, **** p-adjust < 0.0001, ** p-adjust < 0.01, ns not significant)

S1D

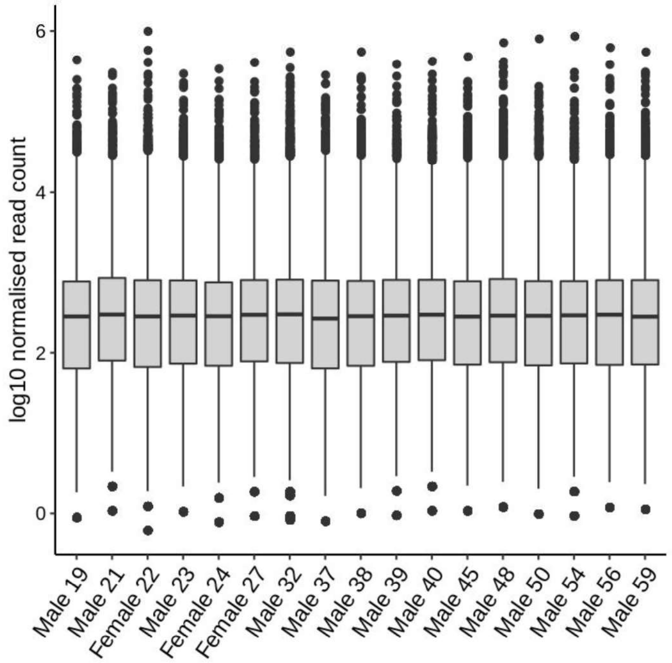


S1E

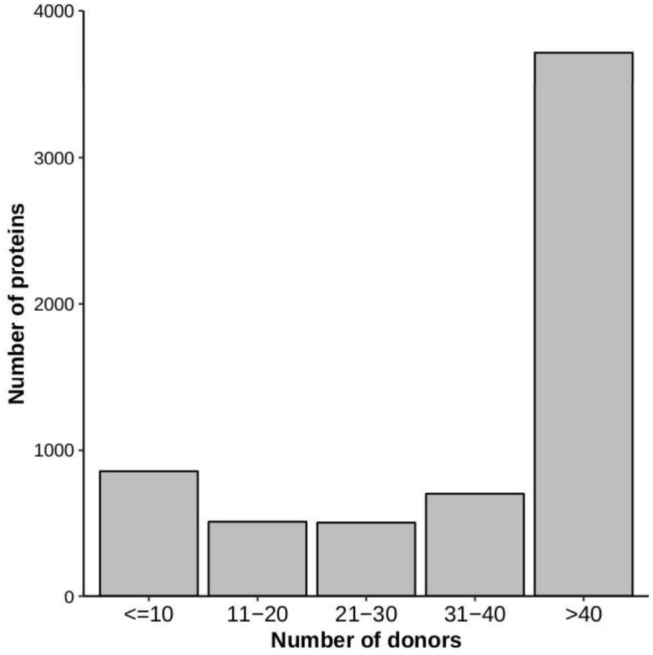


Figure S1D. log10 normalised read count of BMSC RNA-Seq data

Figure S1E. Number of proteins measured in the number of donors

S2A

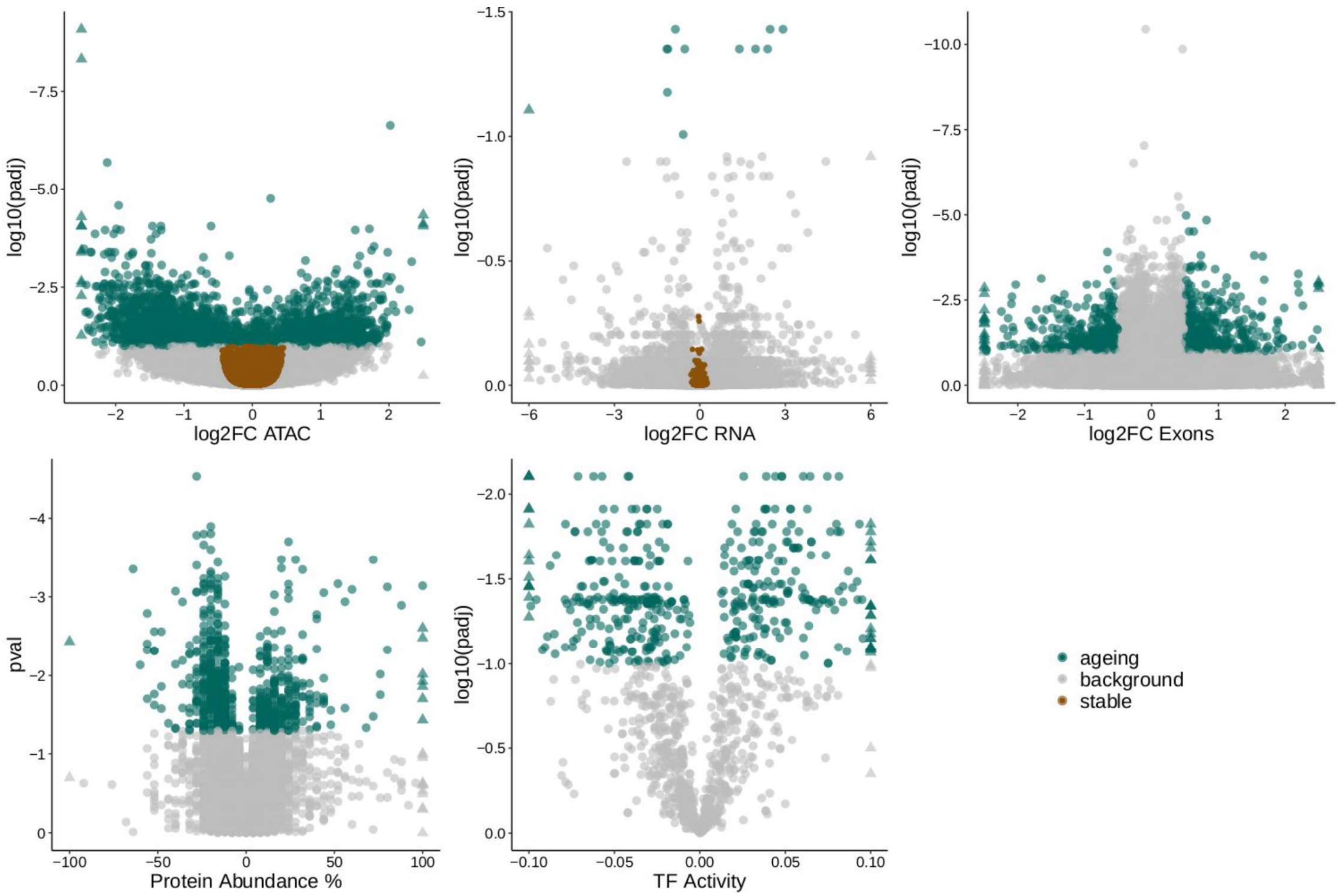


Figure S2A. Age-sensitive elements on the chromatin, RNA, splicing, protein and TF activity levels (green, top left to right, bottom left to right). Stable chromatin regions and transcripts in ageing shown in brown.

S2B

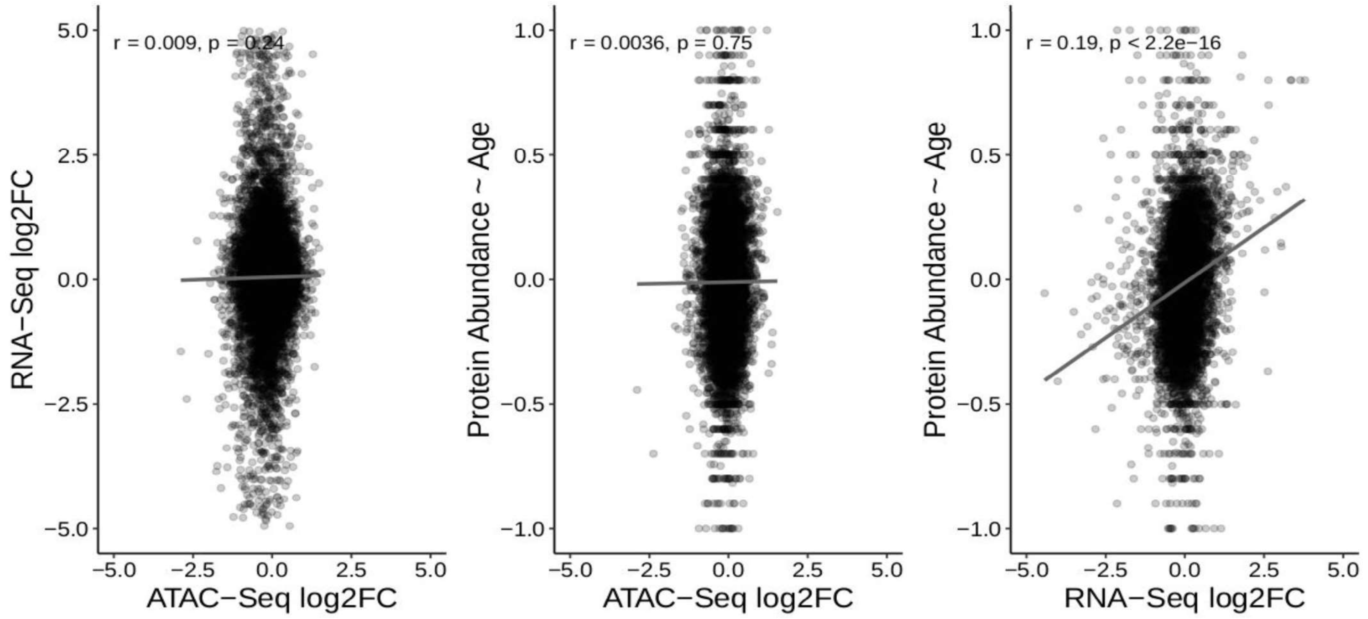


Figure S2B. Correlation of log2 fold change (Old vs Young) and protein abundance in ageing between ATAC-Seq, RNA-Seq and protein data.

S3A

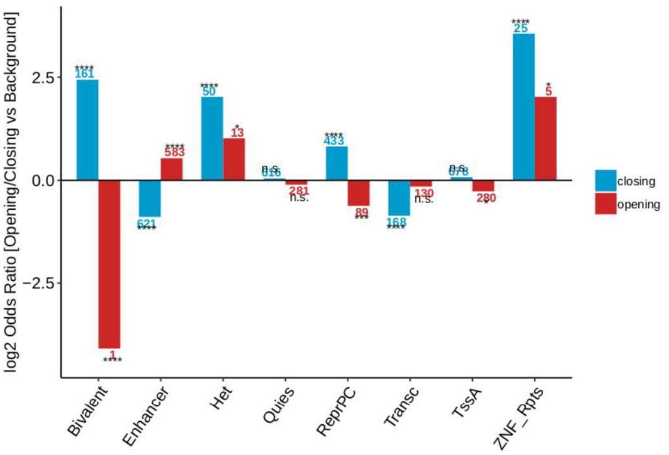


S3B

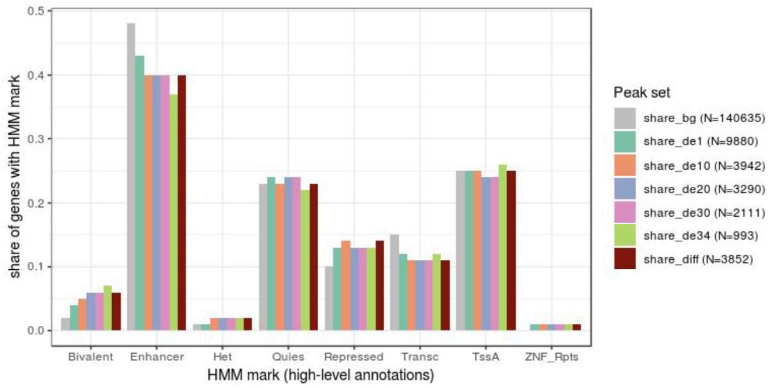


S3C

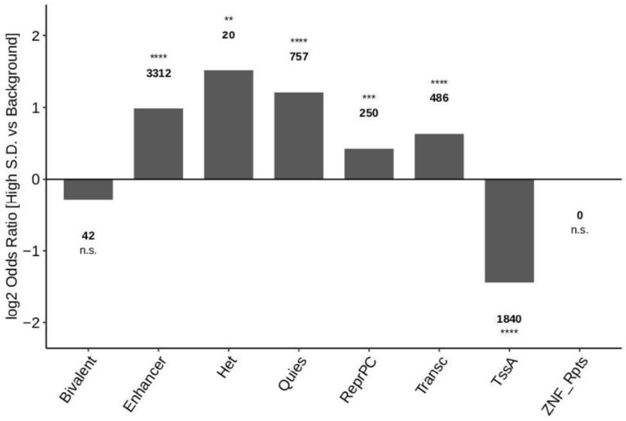


Figure S3A. Enrichment of ATAC-Seq peaks up- and down-regulated in ageing (opening, closing) in different chromHMM regions (*, **, ***, ****, p-adjusted < 0.05, 0.01, 0.001, 0.0001).

Figure S3B. The proportion of ATAC-Seq peaks in different chromHMM regions in different subsetting consensus peaksets. Subsetting consensus peaksets generated by deleted 1 sample at a time. (de1: union of all differential peaks in 34 subsetting rounds, de10 - de34: differential peaks in at least 10 - 34 subsetting rounds, bg: background peaks, diff: original consensus age-sensitive peaks)

Figure S3C. Enrichment of ATAC-Seq peaks with high standard deviation in different chromHMM regions (*, **, ***, ****, p-adjusted < 0.05, 0.01, 0.001, 0.0001).

# S3D

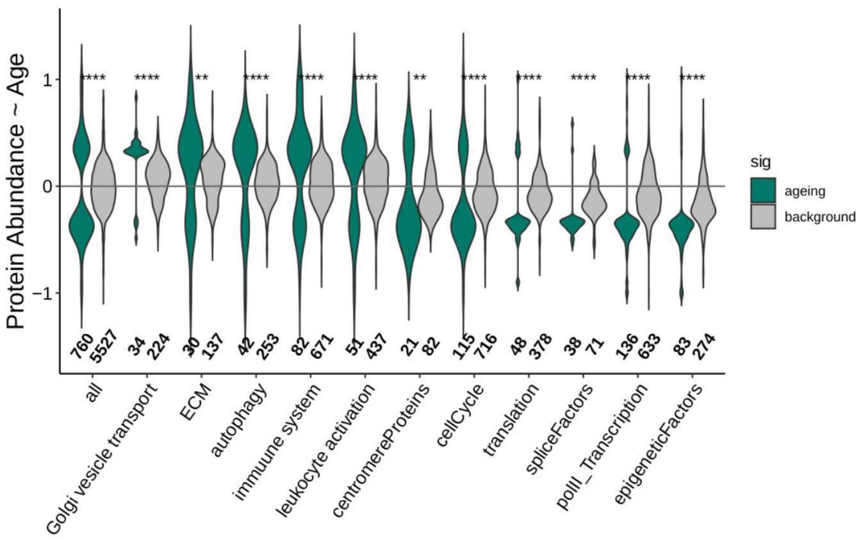


Figure S3D. Distribution of protein changes in different Gene Ontology processes

S3E

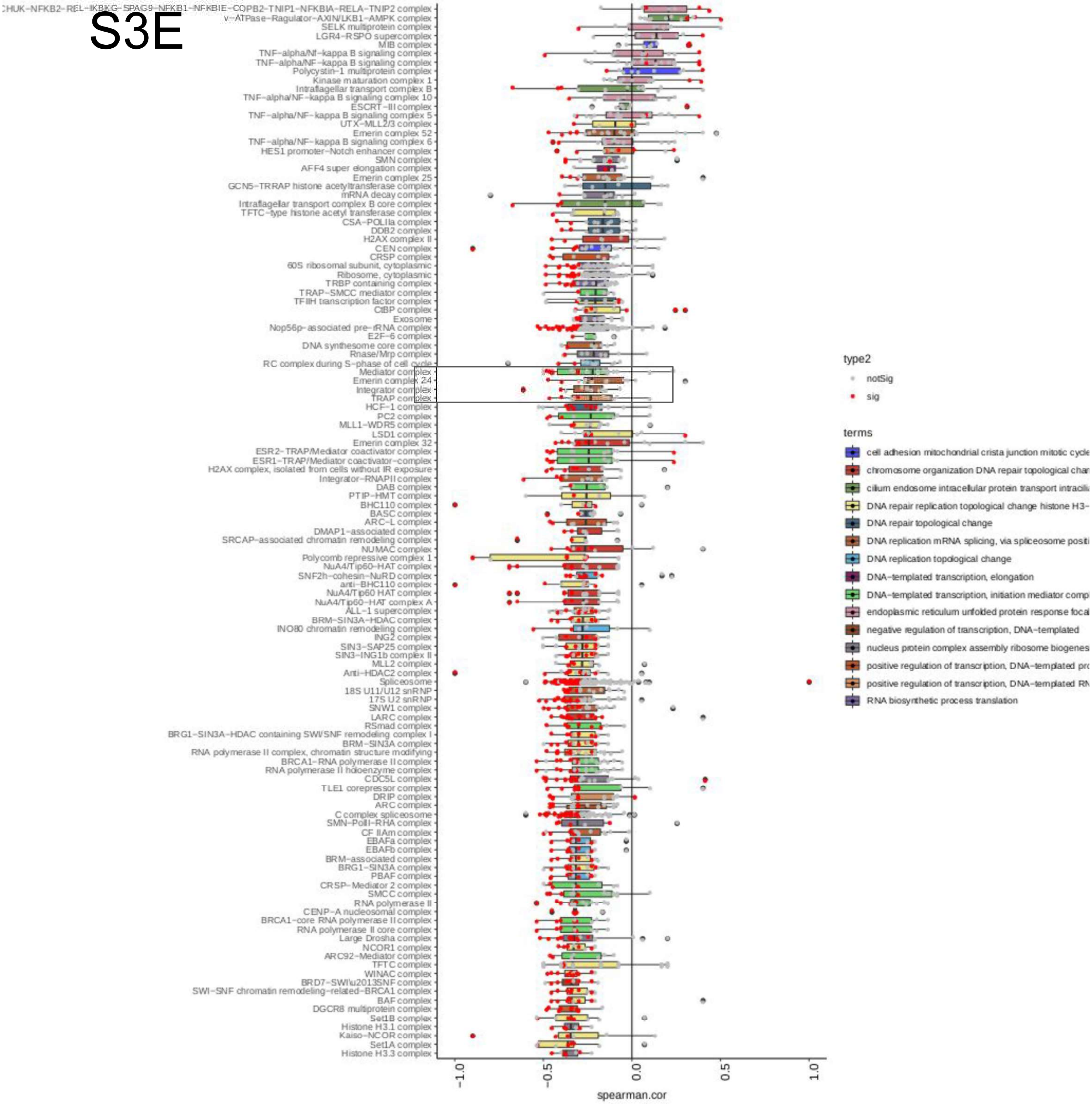


Figure S3E. Proteins abundance vs age (spearman.correlation) in protein complexes. Red represent age-sensitive proteins.

# S3F

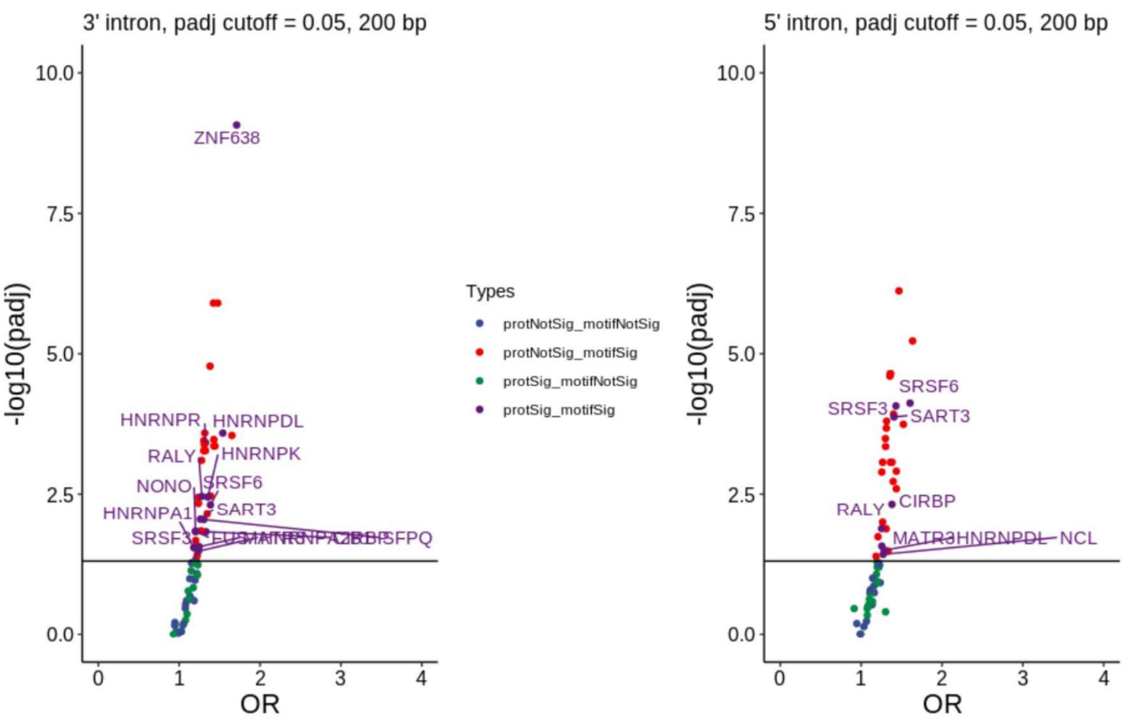


Figure S3F. Enrichment of splicing factor binding sites in 3’ (left) and 5’ (right) intronic regions. Statistical significance of age-sensitive proteins and binding site enrichment indicated (OR: odds ratio).

### Figure S4 A - B

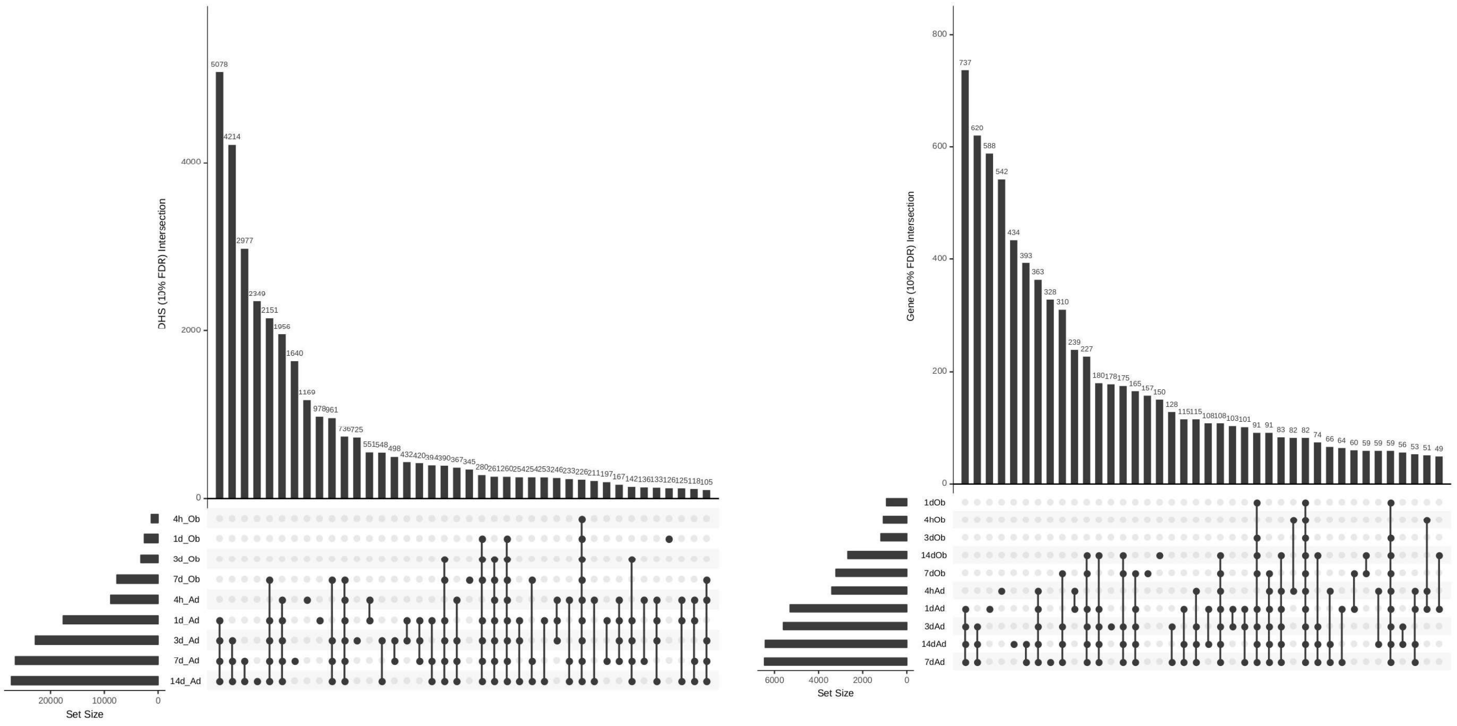


Figure S4. Differentiation elements shared between different differentiation timepoins A) DHS enhancers B) Genes

### Figure S4 C - E

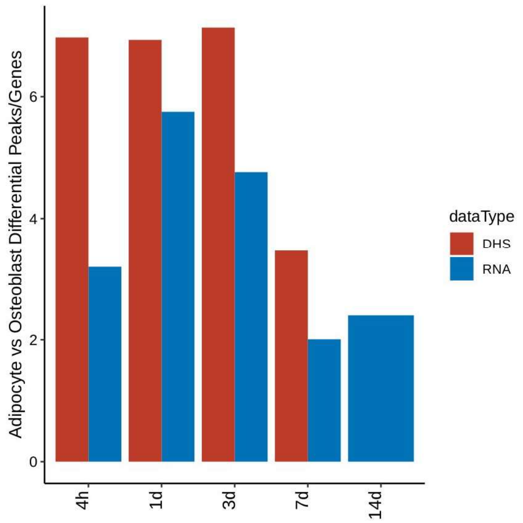


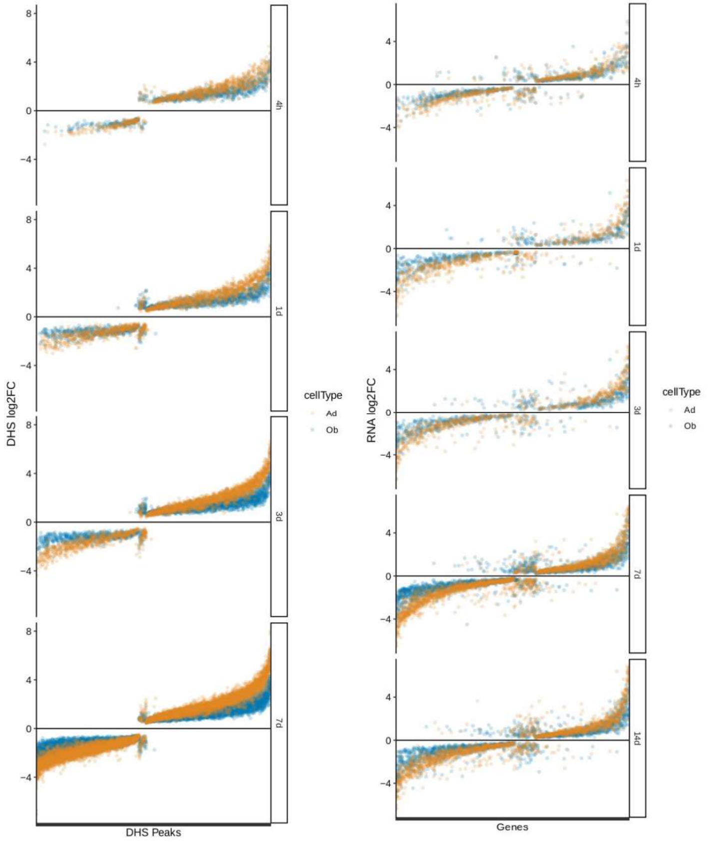


Figure S4A. ratio of differential DHS enhancers and genes in adipogeneis vs osteogenesis.

Figure S4B. log2 fold change of DHS enhancers in adipogenesis (ad) and osteogenesis (os).

Figure S4C. log2 fold change of genes in adipogenesis and osteogenesis.

### Figure S4F

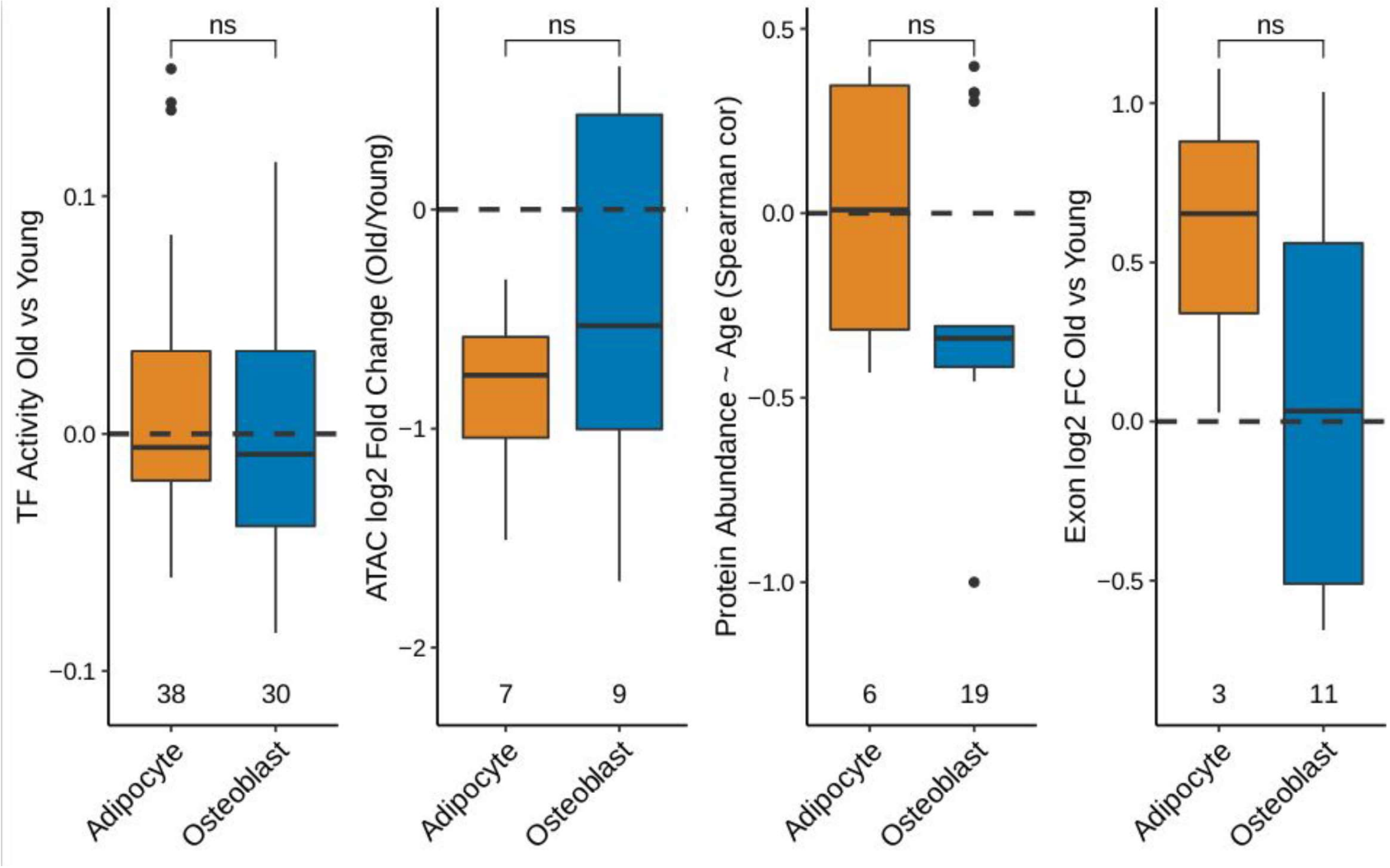


Figure S4F) All TFs, and age-sensitive only ATAC-promoter genes, proteins and differentially used exons, in adipogenesis and osteogenesis GO processes.

### Figure S5

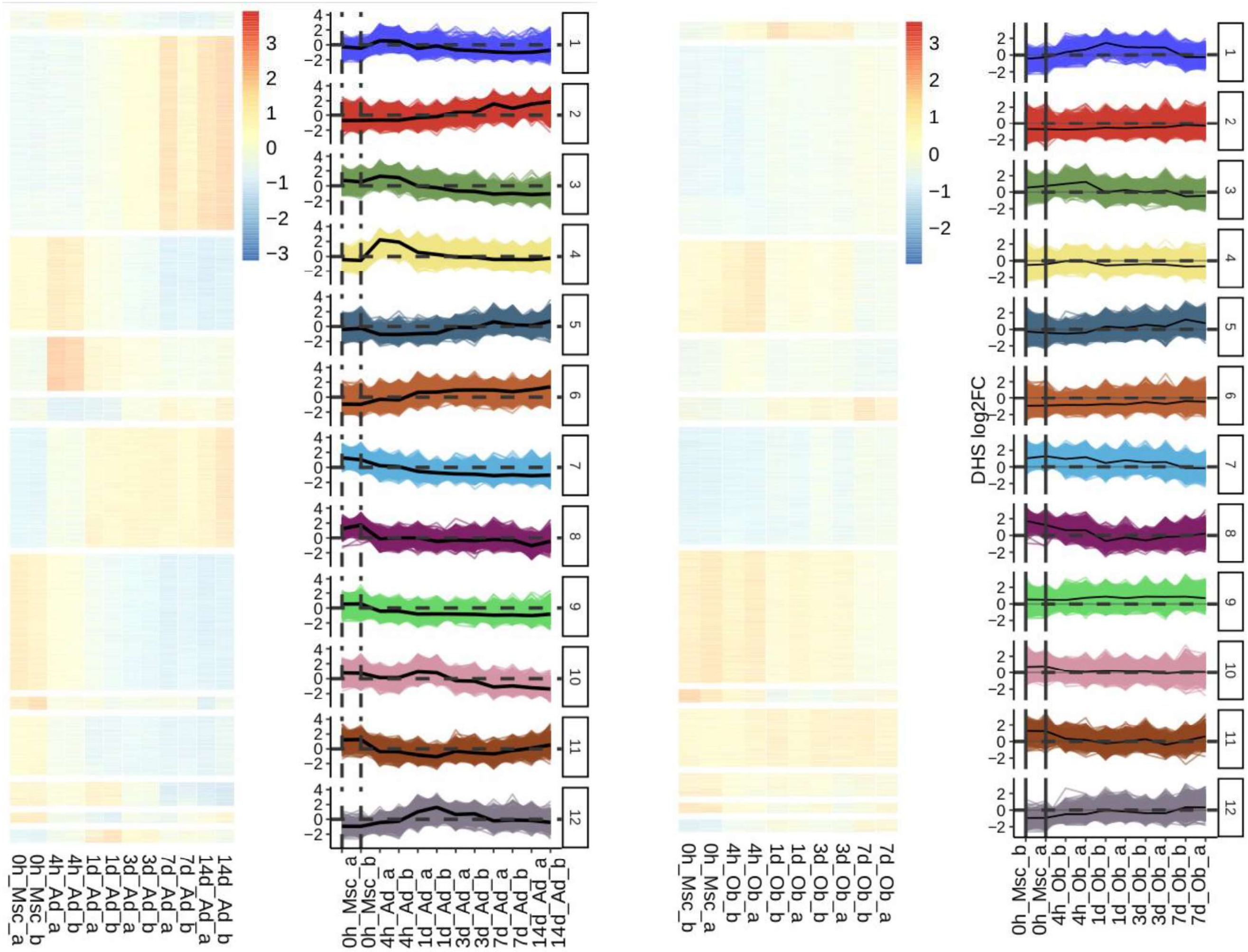


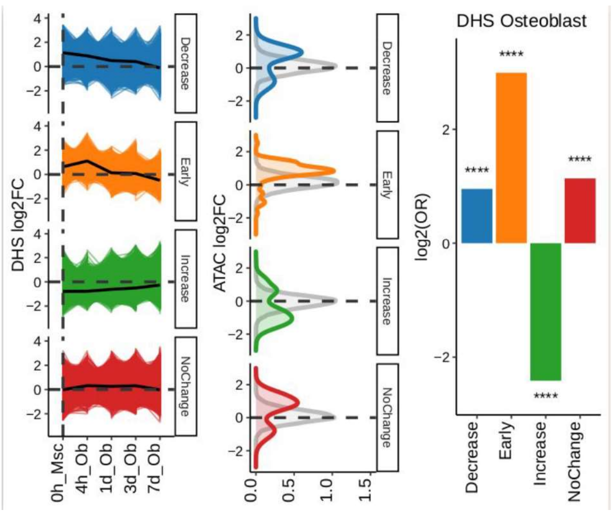


Figure S5. Top panel: Heatmap of z-score of normalised DHS enhancer read counts in adipogenesis and osteogenesis. Peaks were clustered using partition around medoids clustering. The median read counts is summarised in the line plots, which corresponds to the clusters in the heatmap. Clusters classified into broader categories based on their activity throughout the differentiation time course. For adipogenesis: Adipo-ON (clusters 2, 5, 6), Transition-ON (clusters 1, 3, 4), Adipo-OFF (clusters 7, 8, 9, 10), No Change (clusters 11, 12). For osteogensis: Osteo-ON (clusters 2, 4, 5, 12), Transition-ON (clusters 3), Osteo-OFF (clusters 7, 8, 10, 11), No Change (1, 4, 9). Bottom panel: DHS clusters in osteogenesis (log2 fold change differentiated vs undifferentiated), DHS-ATAC overlap (log2 fold change old vs young), enrichment of DHS enhancers in ageing up vs down (Fisher's exact test, **** p-adjust < 0.0001).

### Figure S6

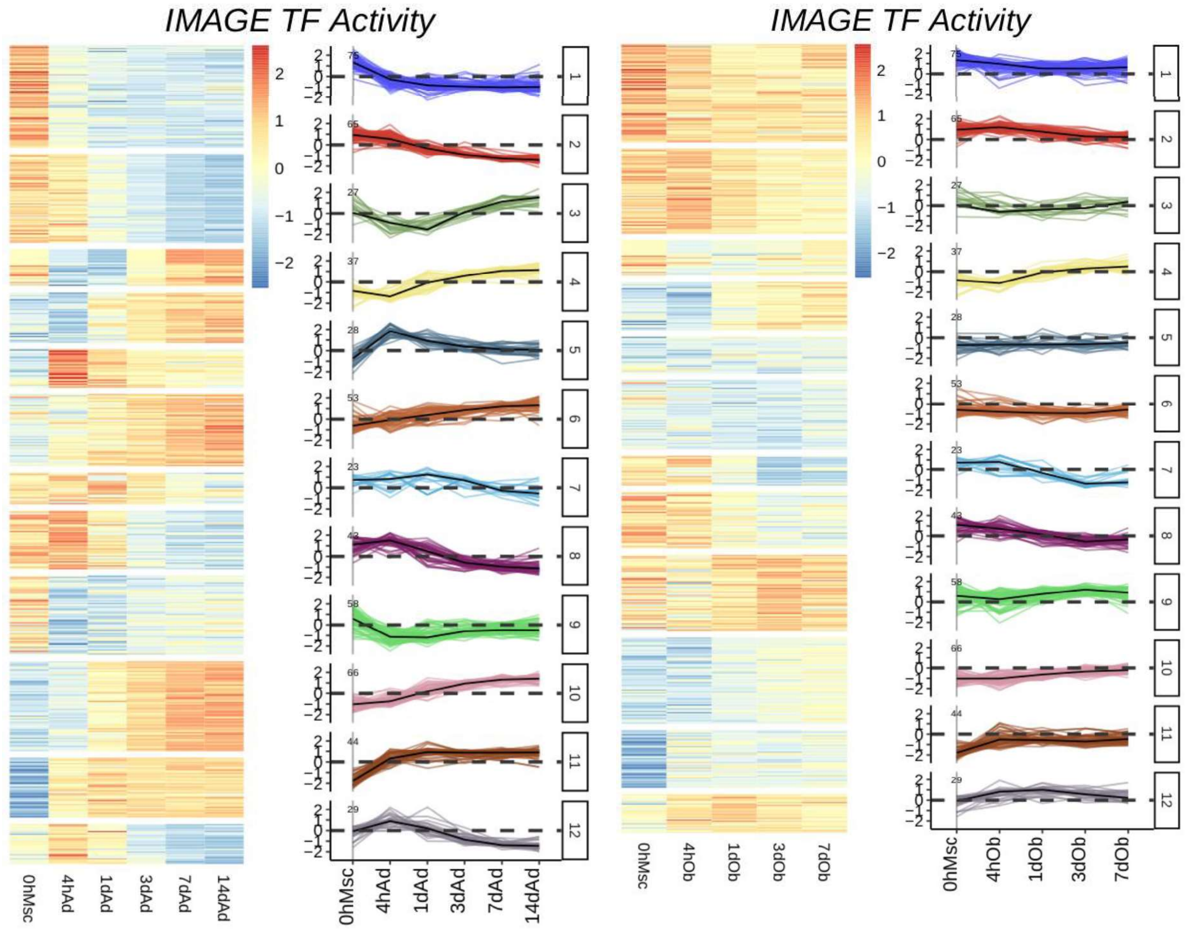


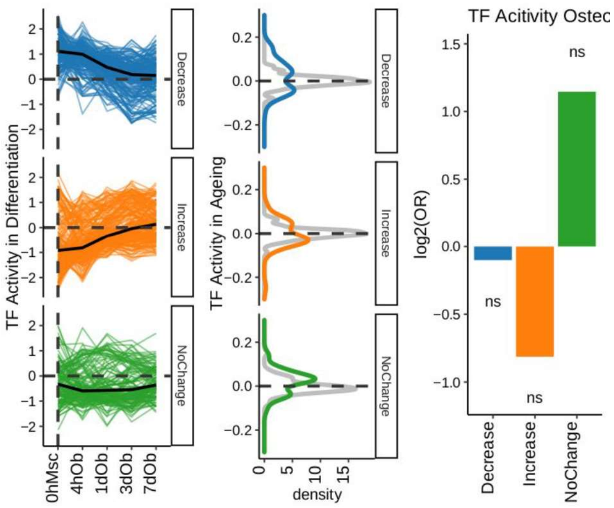


Figure S6. Top panel: Heatmap of z-score of TF activity in adipogenesis and osteogenesis. TFs were clustered using partition around medoids clustering. The median read counts is summarised in the line plots, which corresponds to the clusters in the heatmap. Clusters classified into broader categories based on their activity throughout the differentiation time course. For adipogenesis: Adipo-ON (clusters 3, 4, 6, 10, 11), Transition-ON (clusters 5, 12), Adipo-OFF (clusters 1, 2, 7, 8, 9). For osteogenesis: Osteo-ON (clusters 4, 9, 10, 11), Osteo-OFF (clusters 1, 2, 7, 8), No Change (3, 5, 6, 12). Bottom panel: TF clusters in osteogenesis (log2 fold change differentiated vs undifferentiated), TF overlap with BMSC data (TF activity old vs young), enrichment of TFs in ageing up vs down (Fisher's exact test).

### Figure S7

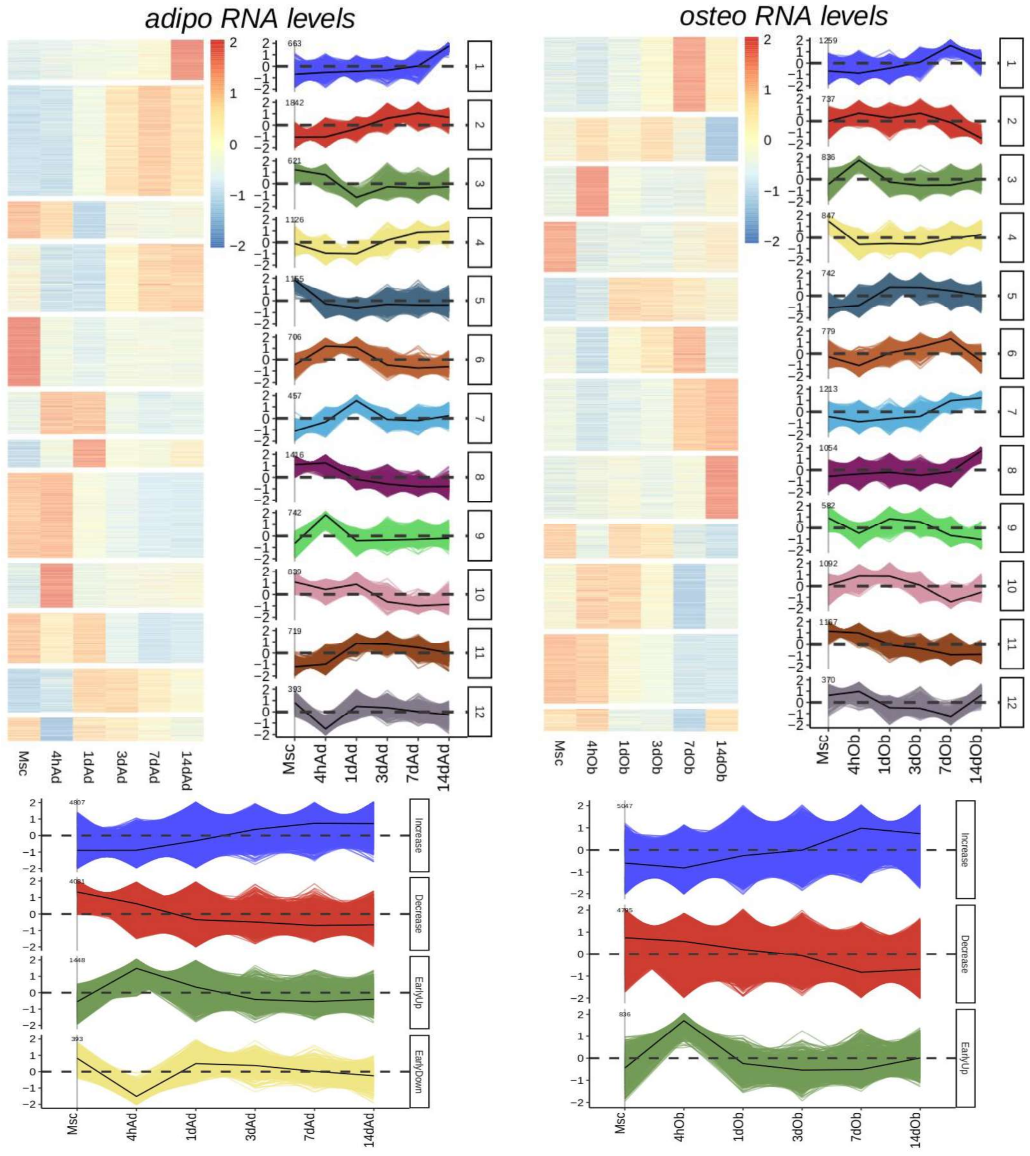


Figure S7. Top panel: Heatmap of z-score of RNA-Seq read counts in adipogenesis and osteogenesis. Genes were clustered using partition around medoids clustering. The median read counts was summarised in the line plots, which corresponds to the clusters in the heatmap. Clusters classified into broader categories based on their activity throughout the differentiation time course. For adipogenesis: Adipo-ON (clusters 1, 2, 4, 7, 11), Transition-ON (clusters 6, 9), Adipo-OFF (clusters 3, 5, 8, 10), Transition-OFF (cluster 12). For osteogenesis: Osteo-ON (clusters 1, 5, 6, 7, 8), Osteo-OFF (clusters 2, 4, 9, 10, 11, 12), Transition-ON (cluster 3). Bottom panel: Gene clusters grouped by activity in adipogenesis (left) and osteogenesis (right).
